# Cholinergic modulation of dentate gyrus processing through dynamic reconfiguration of inhibitory circuits

**DOI:** 10.1101/632497

**Authors:** Mora B. Ogando, Olivia Pedroncini, Noel Federman, Sebastián A. Romano, Luciano A. Brum, Guillermo M. Lanuza, Damian Refojo, Antonia Marin-Burgin

## Abstract

The dentate gyrus (DG) of the hippocampus plays a key role in memory formation and it is known to be modulated by septal projections. By performing electrophysiology and optogenetics we evaluated the role of cholinergic modulation in the processing of afferent inputs in the DG. We showed that mature granule cells (GCs), but not adult-born immature neurons, have increased responses to afferent perforant path stimuli upon cholinergic modulation. This is due to a highly precise reconfiguration of inhibitory circuits, differentially affecting Parvalbumin and Somatostatin interneurons, resulting in a nicotinic-dependent perisomatic disinhibition of GCs. This circuit reorganization provides a mechanism by which mature GCs could escape the strong inhibition they receive, creating a window of opportunity for plasticity. Indeed, coincident activation of perforant path inputs with optogenetic release of acetylcholine produced a long-term potentiated response in GCs, essential for memory formation.

## INTRODUCTION

The dentate gyrus (DG), the main entrance of information to the hippocampus from the entorhinal cortex (EC, (Andersen et al., 1971)), plays a key role in the formation of new memories and memory associations in mammals (Burgess et al., 2002; Leutgeb et al., 2005; Marr, 1971). It is thought to separate the rich repertoire of inputs arriving from EC into non-overlapping sparse patterns of activity, improving discriminability and reducing interference between stored representations (Bakker et al., 2008; Leutgeb et al., 2007; McAvoy et al., 2015; Neunuebel and Knierim, 2014). The sparsity in the activity of granule cells (GCs), the principal neuron type of the DG, is governed by strong local GABAergic inhibition (Nitz and McNaughton, 2004; Pernia-Andrade and Jonas, 2014). In the DG, synaptic inhibition is provided by two main interneuron types, parvalbumin-expressing perisomatic-inhibiting fast-spiking interneurons (PVIs) and somatostatin-expressing dendrite-inhibiting cells (SOMIs) (Freund and Buzsaki, 1996; Hosp et al., 2014; Rudy et al., 2011). PVIs provide powerful feedforward and feedback inhibition to large populations of GCs (Sambandan et al., 2010) and they are critical for the generation of gamma activity (Bartos et al., 2007; Struber et al., 2017). On the other hand, SOMIs (i.e. classically called hilar-perforant-path-associated interneurons-HIPPs) generally distribute axon fibers along the molecular layer, and are engaged by converging GC-inputs providing mostly dendritic feedback inhibition to the DG circuitry, controlling local synaptic and dendritic conductances (although see Yuan et al., 2017 for a detailed characterization of SOMIs subpopulations in the DG).

The GC population is heterogeneous, mostly due to the existence of adult neurogenesis in the DG (van Praag et al., 2002). Although immature GCs receive low excitatory innervation that could limit their recruitment (Dieni et al., 2016; Li et al., 2017), they also receive less inhibition and tend to be more excitable, less selective and more prone to exhibit plastic changes than mature GCs (Danielson et al., 2016; Ge et al., 2007; Marín-Burgin et al., 2012; Mongiat et al., 2009; Pardi et al., 2015). Importantly, mature GCs, despite their sparse activation, reduced excitability and higher threshold for LTP induction, do engage in the formation of memory engrams (Liu et al., 2012; Ramirez et al., 2013; Ryan et al., 2015; Stone et al., 2011; Tronel et al., 2015). In order to be integrated into engrams, mature GCs have to overcome the strong inhibition they receive. While some studies suggested that cell-intrinsic properties could allow them to undergo plasticity (Lopez-Rojas et al., 2016; Lopez-Rojas and Kreutz, 2016), circuital mechanisms could also be involved. One possibility is that neuromodulators adapt circuit processing to allow for plasticity (Palacios-Filardo and Mellor, 2018). The hippocampus receives cholinergic afferents arriving primarily from the medial forebrain (Mesulam et al., 1983; Wainer et al., 1985), that modulate cortical circuit activity (Dannenberg et al., 2017). In particular, acetylcholine (ACh) has a role in hippocampal-related cognitive functions, including coding of spatial location, movement speed, learning and memory (Ballinger et al., 2016; Haam and Yakel, 2017), and attention to sensory stimuli (Bloem et al., 2014; Pinto et al., 2013; Sarter et al., 2005).

Although much is known about the role of ACh as a modulator of information flow at CA1 and CA3 hippocampal subregions (Hasselmo, 2006), cholinergic modulation of the DG microcircuit has been less explored. Recent efforts have been made to explore how septal ACh modulates intrinsic GC firing (Pabst et al., 2016), and its role in memory formation (Raza et al., 2017). However, it remains unknown whether the processing of afferent inputs in the DG microcircuit is changed upon cholinergic modulation. We provide evidence for a cholinergic dynamic reconfiguration of the DG network induced by a nicotinic dependent perisomatic disinhibition of mature GCs, which potentiates responses to afferent inputs. The results presented here shed light on the mechanisms that permit a dynamic adaptation of processing and plasticity upon modulatory states.

## RESULTS

### Cholinergic activation potentiates the response of mature granule cells to afferent inputs

To examine how ACh affects processing of afferent inputs in the DG, we performed electrophysiological loose-patch recordings from mature and immature 4 weeks old GCs (mat-GCs and 4wpi-GCs, respectively) of mouse hippocampal slices. We obtained activation profiles of individual GCs, while electrically stimulating the medial perforant pathway (mPP, Figure 1A). The cholinergic agonist carbamylcholine (carbachol, CBC) was bath applied to study the effect of a high acetylcholine (ACh) neuromodulatory state on the observed activation profiles. We adjusted the intensity of the stimulus delivered to mPP at the threshold of each GC, i.e. an intensity that elicited GC spiking with 50% probability. Five trains of 10 pulses were delivered to the mPP at 1, 10, 20, and 40 Hz for each cell. Adult-born GCs were labeled by injecting a retrovirus in the dorsal DG that led to the expression of RFP or GFP in dividing cells, and acute hippocampal slices were prepared 4 weeks post retroviral injection (4 wpi). As we have previously shown (Pardi et al., 2015), the average number of spikes in response to mPP stimulation in control conditions decreases as stimulation frequency increases (Figure 1B and 1C), with a more drastic effect observed in mat-GCs, indicating that dentate GCs have frequency filters with variable gain depending on their age (Pardi et al., 2015). Nevertheless, CBC induced a strong enhancement of the responses at high frequencies, specifically in mat-GCs, changing the filter properties of this population of GCs. This effect was more variable in 4wpi-GCs and CBC did not significantly change the mean responses of 4wpi-GCs to afferent stimulation (Figure 1C).

**Figure 1.**
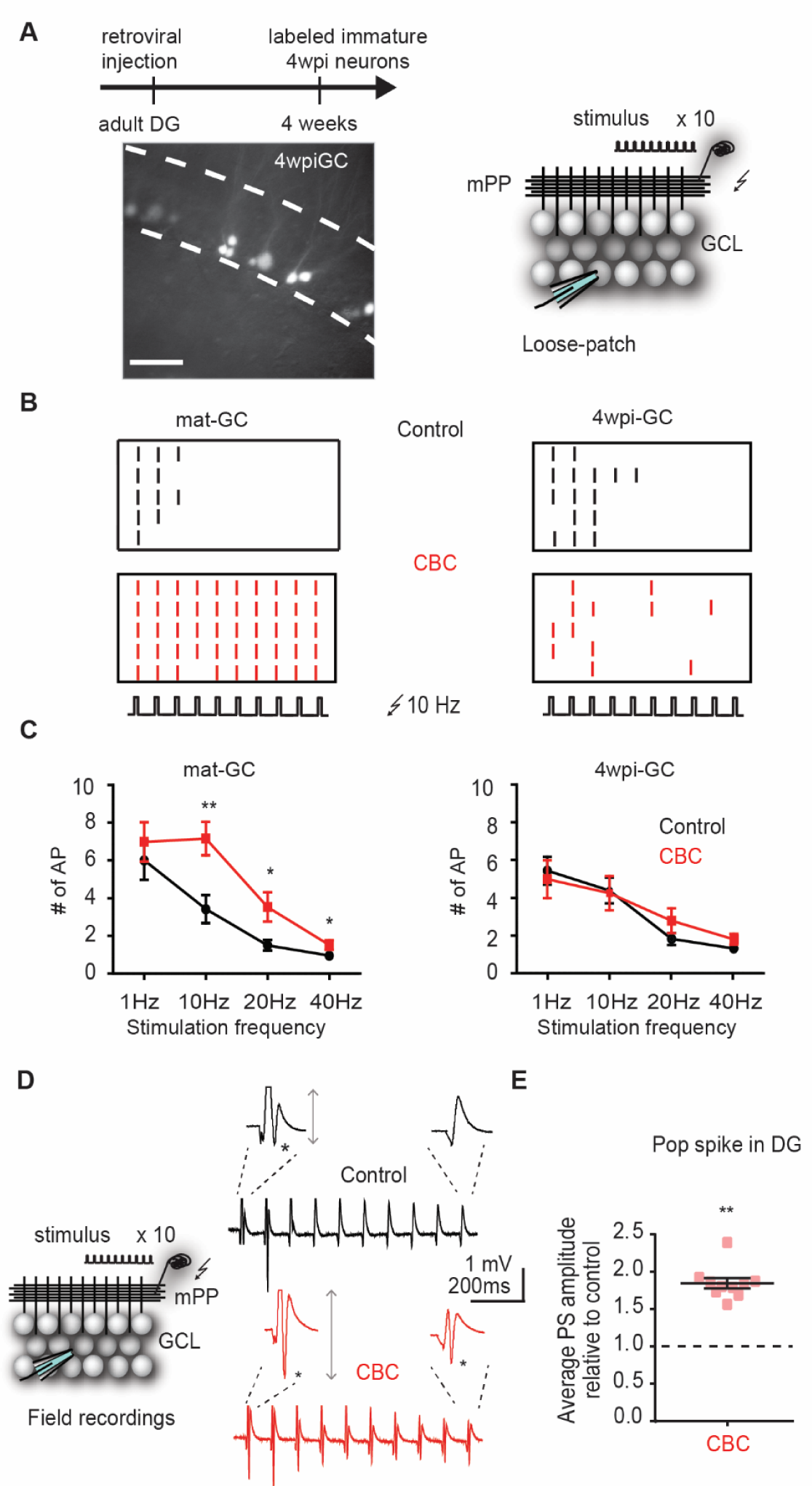
Carbachol increases the spiking response of mature GCs. (A) Experimental design. The image shows a hippocampus slice with 4-weeks-old GC (4wpiGC) expressing RFP. Scale bar: 50 µm. The upper timeline indicates the time of retroviral injection. Right: The scheme shows the recording configuration: a stimulating electrode was placed in the medial perforant path (mPP) to deliver 10 stimuli at different frequencies; the stimulation intensity was kept at 50% spiking probability for each neuron. Loose-patch recordings were obtained from mature GCs (mat-GC) and 4wpi-GCs to detect spikes. (B) Raster plots from one mat-GC (left) and one 4wpi-GC (right) at 10 Hz in the control condition (upper panels, black) and after bath application of CBC (lower panels, red). Each color dash denotes a spike. (C) Average sum of action potentials evoked by stimulation trains of 1 Hz, 10 Hz, 20 Hz, and 40 Hz, in mat-GCs (left) and 4wpi-GCs (right) in the control condition (black) or after bath perfusion with CBC (red). CBC increased spiking response of mat-GC but not 4wpi-GC. (mat-GC: two-way ANOVA, variation between control and CBC, F_1, 134_= 21.61: \*\*\**p* < 0.0001; variation in frequency F _3, 134_= 23.44: \*\*\**p < 0.001*; interaction F _3, 134_ = 2.800: \**p =* 0.0425; N≥7 cells from 5 mice. 4wpi-GC: two-way ANOVA, variation between control and CBC, F _3, 112_ = 0.5046: ns, *p = 0.6799*; variation in frequency, F _3, 112_ = 11.69: \*\*\**p* < 0.001; interaction F_1, 112_ = 0.02754: ns, *p =* 0.8685; N≥9 cells from 5 mice). (D) The scheme shows the recording configuration: A stimulating electrode was placed in the mPP to deliver 10 stimuli at 10 Hz while an extracellular recording electrode was placed in the granule cell layer (GCL) to assess population spike (PS) responses. Right: Example traces showing the population spike response to a train of 10 pulses at 10 Hz before (control, upper black) or after bath perfusion with CBC (CBC, bottom red). (E) Average amplitude of the population spike response to the 10 Hz train after bath application of CBC (values are relative to control conditions, red). CBC increased the population spike responses in the DG (paired t-test, control vs CBC *p < 0.05, N=7 slices from 3 mice).

As mature neurons represent more than 95% of the total population of GCs in the adult DG, we hypothesized that the effect of CBC would also impact on the collective spiking of the GC population. Because the biggest effect in individual neurons was observed upon 10 Hz stimulation of the mPP, we studied the mechanisms of action of the cholinergic modulation at this frequency. This frequency is physiologically relevant, since the DG predominantly oscillates at frequencies near 10 Hz in vivo (Jung and McNaughton, 1993; Leutgeb et al., 2007; Neunuebel and Knierim, 2012), and the medial septum is involved in the generation of these oscillations (Carpenter et al., 2017; Kang et al., 2017).

Population spike (PS) responses were obtained by placing an extracellular electrode in the GCL of the DG and stimulating the mPP fibers with trains of 10 pulses at 10 Hz (Figure 1D). Consistent with loose-patch recordings, population spike amplitude in control conditions decreased strongly along the pulses of the stimulation train, as there are less GCs that respond at the last pulses (Figure 1D). As expected, bath application of CBC produced an increase in the population spike responses to afferent stimuli at 10 Hz in the GCL (Figure 1E). The increase in GC activation is more prominent in the last pulses of the train, but also significant in the first pulses (average gain in activation was 1.86 ± 0.44; gain in activation in the first 5 pulses 1.33 ± 0.34; gain in activation in the last 5 pulses 3.18 ± 0.46). Importantly, the increase in GCs activity is transferred postsynaptically since we observed a concomitant significant increase in the evoked activity of CA3 (Figure S1 D-F).

To better control the spatial and temporal release of ACh, we explored if endogenous release of ACh from septal axons would produce a similar increase in the activity of the DG. For this, we used the double transgenic mouse line ChAT-ChR2, which expresses the photoactivatable protein channelrhodopsin-2 (ChR2) in cholinergic neurons of the septum and in the axons that project and innervate the DG (Figure 2A) (Dannenberg et al., 2019; Herman et al., 2016; Yaeger et al., 2019). We obtained hippocampal slices from these animals and elicited endogenous release of ACh from cholinergic terminals by stimulating with blue light (Figure 2B). We recorded population spikes in response to mPP stimulation before and after the blue light illumination protocol (5-seconds trains of light pulses at 10 Hz delivered every minute, paired with electrical activation of mPP, Figure S2A-C). Endogenous ACh release also augmented the population spike responses in the DG upon 10 Hz electrical stimulation of mPP compared to the control condition before illumination (Figure 2C). The average gain in activation was 1.33 ± 0.06, and it was more prominent at the initial pulses (1.57 ± 0.14), but also significant for the last pulses of the train (Figure 2D). Importantly, evoked responses remained constant during the same optogenetic stimulation protocol in Cre ^-/-^ animals (Figure 2E), as well as in Chat-Tom animals (Figure S2 D). The modulation of GCs responses by ACh was not an ON/OFF phenomenon but outlasted light stimulation of ACh fibers (23.0 ± 6.7% 10 min after light off, *p<0.05, paired t-test, N = 6, Figure S2 D). This neuromodulatory gain, thus, allows GCs to respond more strongly and to follow a wider frequency range of afferent stimuli.

**Figure 2.**
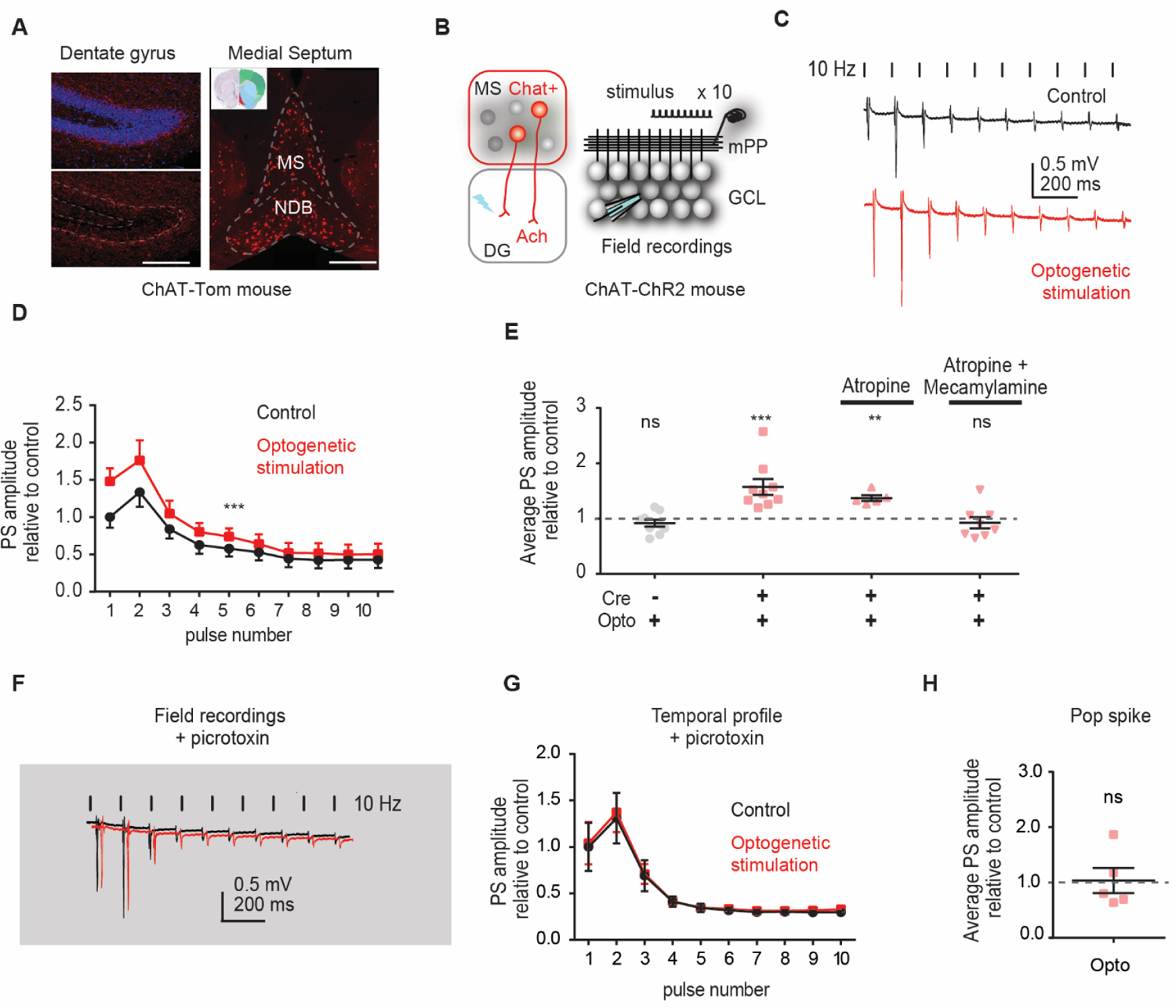
Endogenous release of acetylcholine increases spiking response in the DG through inhibitory circuits. (A) Experimental design. The left image shows a hippocampal slice obtained from adult ChAT-Tom mice with cholinergic terminals expressing tomato (red), but no somas, in the DG. Somas are labeled with DAPI in the upper panel. The right image shows an anterior brain slice obtained from ChAT-Tom mice where the medial septum (MS) and the diagonal band of broca (DB) contain tomato expressing neuronal somas. Scale bar: 500 µm (B) Experimental scheme, in slices obtained from ChAT-ChR2 mice, we use blue light to activate the cholinergic terminals in the DG. Right, recording configuration: a stimulating electrode was placed in the medial perforant path (mPP) to deliver 10 stimuli at 10 Hz; Extracellular recordings were obtained from the GCL to detect pop-spikes (PS). (C) Example traces showing the population spike response to a train of 10 pulses at 10 Hz before (Control, upper black) or after optogenetic stimulation protocol to activate cholinergic terminals (red) (see Figure S2). (D) Average amplitude of the population spike at each pulse (relative to the 1^st^ pulse in control condition) of the 10 Hz train in control (black) and after optogenetic stimulation protocol (red). Optogenetic stimulation increased PS response in the DG (two-way ANOVA, variation between control and optogenetic stimulated F _1, 80_ = 133.9: \*\*\**p* < 0.0001; variation between pulses F _9, 80_ = 7.152: \*\*\**p*< 0.0001; interaction F _9, 80_ = 8.185: \*\*\**p* < 0.001, N=9 slices from 5 mice). (E) Average amplitude of the population spike response to the 10 Hz train in control condition (black) and after light stimulation protocol (red, Opto) with or without the cholinergic antagonists atropine and mecamylamine. Bath perfusion with a combination of atropine and mecamylamine prevented optogenetic induced potentiation of population spike responses (One sample t-test Cre ^-/-^, Control vs Opto p>0.05 N=9 slices from 3 mice; Cre ^+/-^, Control vs Opto **p < 0.01, N=9 slices from 5 mice; Cre ^+/-^, Control vs Opto Atropine **p < 0.01, N=5 slices from 3 mice; Cre ^+/-^, Control vs Opto Atropine + Mecamylamine ns p >0.05, N=8 slices from 3 mice). (F) Example traces showing the population spike response to a train of 10 pulses at 10 Hz before (Control, black) or after optogenetic stimulation protocol (Opto, red) to activate cholinergic terminals. Experiments were performed in the presence of the GABA-A antagonist picrotoxin (PTX). (G) Pop-spikés average amplitude for each pulse of the 10 Hz train relative to the 1^st^ pulse in the control condition (black) and after optogenetic stimulation (red), in the presence of PTX. (H) Mean population spike amplitude in response to the 10 Hz train in control condition (black) and after light stimulation protocol (red, Opto). Light stimulation did not change the population spike response of GCs in presence of PTX (two-way ANOVA, variation between control and light stimulated F _1, 8_ = 0.04888: *p* > 0.9999; variation between pulses F _9, 72_ = 24.51: \*\*\**p*<0.0001; interaction F _9, 72_ = 0.01556 ns *p* = 0.8306, N=5 cells from 3 mice).

To confirm that this cholinergic modulation was due to activation of septal cholinergic terminals, we performed stereotaxic injections of an adenovirus carrying Cre-dependent ChR2 expression (AAV9-CAG-DIO-ChR2) delivered to the medial septum of ChAT^Cre^ mice. In slices obtained from these animals, pairing optogenetic activation of septal cholinergic terminals with afferent activation also induced an increase in the population spike response (Figure S3).

In order to verify that the modulation of mat-GCs’ activity was indeed cholinergic, we pharmacologically blocked ACh receptors with different antagonists. We used broad spectrum antagonists for both muscarinic (atropine) and nicotinic (mecamylamine) receptors. As expected for a cholinergic modulation, perfusion with ACSF containing these antagonists prevented the optogenetically induced potentiation of GC responses (Figure 2E). Interestingly, bath application of atropine alone was insufficient to block the cholinergic neuromodulatory effects, suggesting that nicotinic receptors are primarily involved in the signaling pathway (Figure 2E).

Activation of mPP not only generates excitation but it also recruits inhibitory circuits that regulate the activation of GCs (Marín-Burgin et al., 2012). We hypothesized that the increase in the amplitude of the population spike upon cholinergic activation could be generated by an increase in the excitatory inputs, a decrease in the inhibitory inputs or a change in the intrinsic properties of the neurons. To explore the underlying mechanism, we first evaluated the involvement of inhibitory circuits by pharmacologically blocking GABA_A_ receptors with picrotoxin (Figure 2F). In this condition, optogenetic activation of cholinergic terminals in the DG did not induce a potentiation of the population spike response, indicating that cholinergic neuromodulation requires intact inhibitory circuits (Figure 2F-H). This result also suggests that cholinergic modulation does not involve a change in excitation.

Last, to evaluate if the effect of cholinergic modulation was due to a change in the intrinsic properties of the neurons, we measured the resting membrane potential (Vm), the action potential threshold and the input resistance (IR) of neurons upon activation of cholinergic inputs. Although CBC induced a small depolarization of the neuronal membrane potential as previously reported (Muller et al., 1988), the endogenous release of ACh did not affect Vm, the threshold or the IR of the recorded neurons (RMP-76.53 ± 2.03 vs −76.13 ± 3.37; IR 522.3 ± 58.46 vs 579.4 ± 42.59; threshold −40.2 ± 2.05 vs −42.2 ± 1.67, ns paired t-test, N=7 cells from 3 mice), further suggesting that the increased responses of neurons upon cholinergic modulation is not explained by a modification of the intrinsic cell properties, but presumably due to a modulation of the inhibitory circuits.

### Synaptic mechanisms underlying cholinergic modulation of granule cells

Because blocking inhibitory circuits prevents the cholinergic-induced increase in activity of GCs, we investigated the precise contribution of the evoked excitatory and inhibitory inputs to GCs at different modulatory states. We performed whole-cell voltage-clamp recordings of mat-GCs to measure excitatory and inhibitory responses elicited at 10 Hz stimulation of mPP axons in hippocampal slices from ChAT-ChR2 mice (Figure 3A). In this configuration, we isolated inhibitory postsynaptic currents (IPSCs) and excitatory postsynaptic currents (EPSCs) of each neuron by clamping the voltage of the neuron at the reversal potential of excitation (∼0 mV) or inhibition (∼-60 mV), respectively (Figure 3B). Importantly, direct monosynaptic IPSC elicited in presence of kynurenic acid (KYN) was subtracted from the total IPSC at the end of each recording for the analysis. As expected, combining optogenetic release of endogenous ACh with activation of mPP induced a slow but strong decrease in the evoked IPSC (Figure 3C) that outlast the optogenetic stimulation (Figure S2), whereas EPSC was largely unaffected by cholinergic activation (Figure 3D). This produced an increase in the E/I balance of neurons that explains the changes in the activity profile (Figure 3E). Because different ACh receptors are known to be expressed in both GCs and interneurons, the overall effect could be a combination of multiple pathways activated in different cell types. In agreement with our results at the population activity level, bath application of the nonspecific nicotinic antagonist mecamylamine prevented the light-evoked reduction in the IPSC (Figure 3F, Figure S2F), confirming the crucial role of nicotinic receptors in the cholinergic modulation of the DG activity, and discarding again unspecific effects of light and/or heat (Allen et al., 2015).

**Figure 3.**
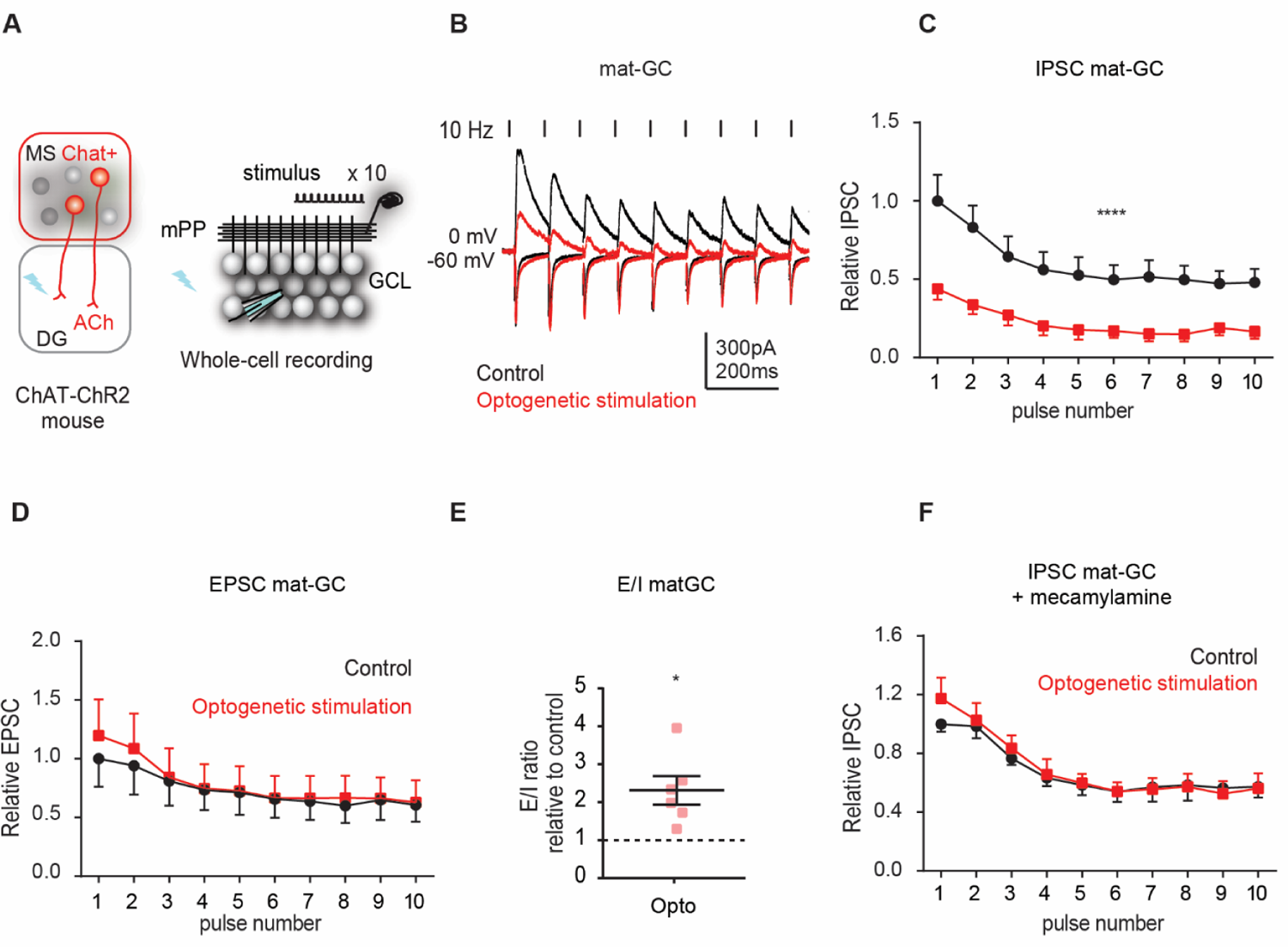
Cholinergic activation decreases evoked inhibitory postsynaptic current (IPSC) onto mat-GCs through nicotinic receptor activation. (A) Experimental scheme. Hippocampal slices were obtained from ChAT-ChR2 mice. Blue light stimulates cholinergic terminals in the DG. Right, recording configuration: A stimulating electrode was placed in the mPP to deliver a train of 10 stimuli at 10 Hz, while whole-cell voltage-clamp recordings were obtained from GCs. Stimulation of the mPP elicits monosynaptic excitation (EPSC) that was recorded at the reversal potential of inhibition (−60mV) and multisynaptic inhibition (IPSC) that was recorded at the reversal potential of excitation (0mV). (B) Example traces of the IPSCs and EPSCs obtained upon mPP stimulation in control (black) or after blue light stimulation protocol to activate cholinergic terminals (Opto, red). The dashed line shows the time of each stimulus. (C) Relative IPSC peak at each pulse in control (black) or after light protocol stimulation (red). Activation of cholinergic axons strongly reduced evoked IPSCs on mat-GCs (Two-way ANOVA, control vs light stimulated F _1, 5_ = 21.39: ** *p* = 0.0057; variation in pulse number F _9, 45_ = 10.42: \*\*\*\**p*<0.0001; interaction F _9, 45_= 9.291: \*\*\*\**p*<0.0001, N=6 cells from 4 mice). (D) Relative EPSC peak at each pulse in control (black) or after light protocol stimulation (Opto, red). Activation of cholinergic axons did not change evoked EPSCs on mat-GCs (Two-way ANOVA, control vs light stimulated F _1, 10_ = 4.606e-007: *p* = 0.9920; variation in pulse number F _9, 90_ = 18.15: \*\*\**p*<0.001; interaction F_9, 90_ = 0.2123: *p* = 0.9995, N=6 cells from 4 mice). (E) Average excitation/inhibition (E/I) balance relative to control elicited by the 10 Hz train after light stimulation protocol (red)). Activation of cholinergic axons increases the E/I balance on mat-GCs (paired t-test, control vs light stimulated: \**p*<0.05, N=6 cells from 4 mice) (F) Relative IPSC peak at each pulse in control (black) or after light protocol stimulation (Opto, red). Bath application of Mecamylamine prevented cholinergic induced reduction of evoked IPSCs on mat-GCs (Two-way ANOVA, control vs light stimulated F _1, 6_ = 0.06525: *p* = 0.8069; variation in pulse number F _9, 54_ = 30.59: \*\*\*\**p*<0.0001; interaction F _9, 54_ = 0.6902: *p* = 0.7145, N=4 cells from 3 mice).

Taken together, these results demonstrate that activation of cholinergic terminals in the DG selectively suppresses inhibitory inputs onto mat-GCs recruited by activation of mPP. Thus, the resulting increase in the E/I balance of these neurons can explain their stronger spiking response to afferent inputs. In fact, the timing of IPSC reduction due to cholinergic release follows the population spike enhancement with a remarkable similar time course (Figure S2D-E), indicating again that the reduction of inhibition is responsible for the increased output of GCs.

### Acetylcholine reconfigures inhibitory circuits

Because ACh release produces a very strong effect on evoked inhibition onto mat-GCs, we further explored which specific component of this inhibition is affected. As previously mentioned, activation of mPP not only produces excitation on GCs but also activates inhibitory circuits in a feedforward manner (Marín-Burgin et al., 2012). In addition, activation of GCs can also recruit feedback inhibition onto themselves and others, and this inhibition can influence their activity, especially during trains of stimuli (Roux and Buzsaki, 2015). These two types of inhibition are known to exert different functions in shaping the firing patterns of principal neurons. The reduction of the mPP evoked IPSC could be due to a reduction of the feedforward component of inhibition or in the feedback component (or both). To assess the feedback component of the IPSC, we placed the stimulating electrode directly in the GCL, electrically activating GCs that recruit feedback inhibition (FB-IPSC) onto other GCs, and thus largely bypassing the mPP axonal activation (Figure 4A). Surprisingly, the FB-enriched component is unaffected by cholinergic modulation (Figure 4B-C, Figure S2F). These results suggest that ACh can specifically decrease the feedforward inhibitory component recruited onto GCs upon mPP activation.

**Figure 4.**
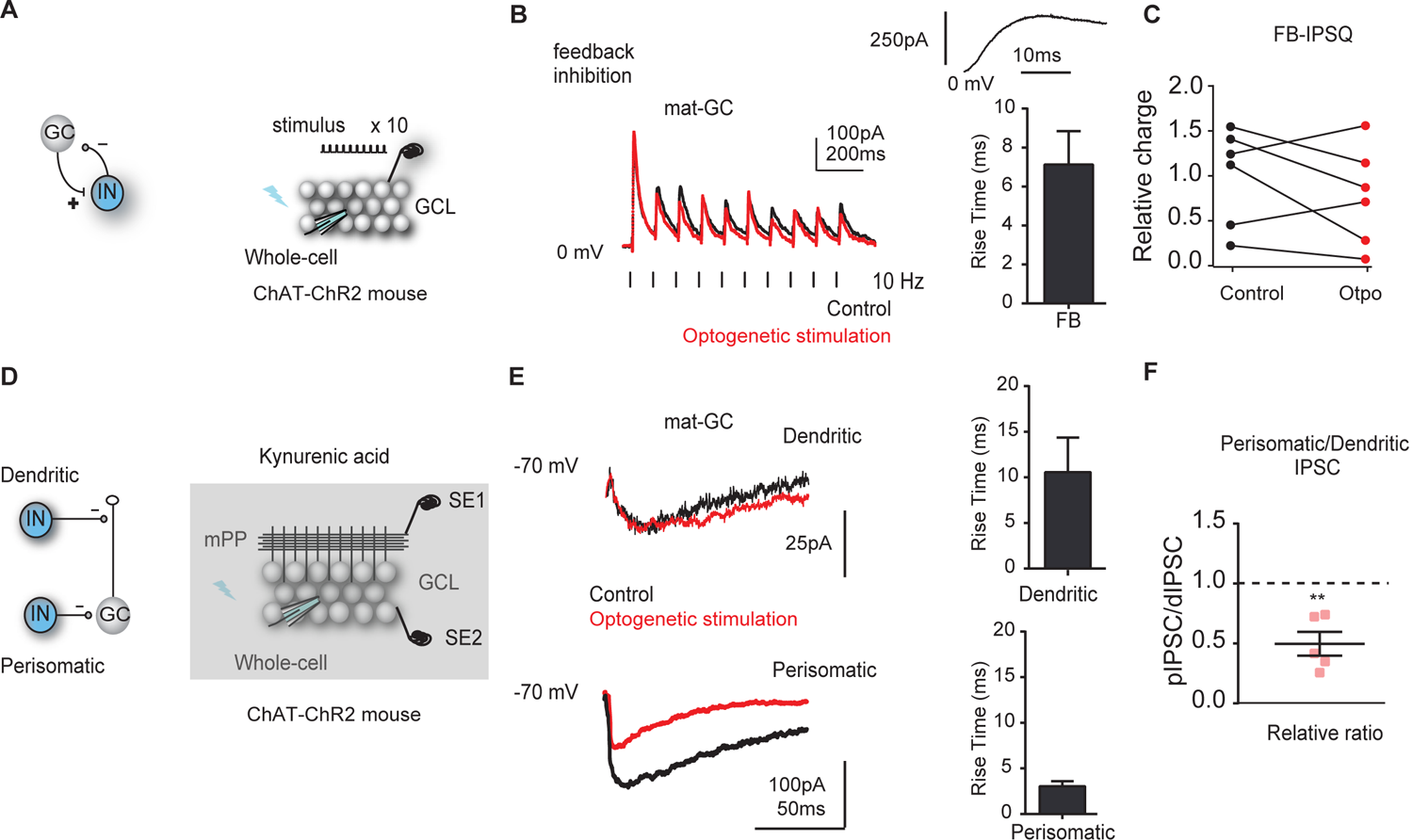
Acetylcholine reduces mPP but not GCL evoked IPSC in mat-GCs. (A) Recording configuration showing the activated circuit. Direct stimulation of the GC activated FBI. The stimulation electrode was placed in the GCL to reduce recruitment of FFI. (B) Example trace of evoked FB-IPSC after GCL stimulation with a train of 10 pulses at 10 Hz in control (black), or after blue light stimulation protocol (red). Right, mean value of the FB-IPSC rise time (10-90%) from the recorded neurons. Upper inset shows the evoked FB-IPSC in an increased scale. (C) Total IPSQ evoked by the train. Light stimulation did not change the FB-IPSQ (control vs light stimulated: *p*>0.05, paired t-test, N=6 from 3 mice). (D) Schematic diagram of the recording configuration. Stimulating electrodes were placed in the GCL and the ML to activate monosynaptic perisomatic or dendritic inhibition. Experiments were performed in the presence of 6 mM kynurenic acid (Kyn) to block glutamatergic transmission. IPSCs were subsequently measured in whole-cell recordings from individual mat-GCs. Perisomatic IPSCs display a reversal potential of ∼0 mV due to the recording conditions under symmetrical Cl^-^, while dendritic IPSCs display a positive driving force at that potential. (E) Example traces of evoked perisomatic (bottom) and dendritic (top) IPSCs from mat-GCs recoded at −70mV in control conditions (black) or after optogenetic stimulation of cholinergic afferents (red). Right, mean value of the perisomatic-IPSC (bottom) and dendritic-IPSC (top) slopes from the recorded neurons. (F) Optogenetic stimulation produced a decrease in the ratio between perisomatic and dendritic IPSCs relative to control (paired t-test, control vs light stimulated: \*\**p* < 0.01, N=5 cells from 3 mice).

To further study the effect of ACh in feedforward-enriched IPSC (FF-IPSC), we pharmacologically reduced the output of the DG GCs by bath application of DCGIV (Figure S4), an agonist of group II metabotropic glutamate receptors (mGluR_2/3_) that reduces release probability in mossy fiber terminals (Kamiya et al., 1996). Application of DCGIV largely reduces feedback inhibition (66.3 ± 7.31 % reduction of FB-IPSC, vs 41.6 ± 4.42 % reduction of total IPSC, Figure S4D-F), and the remaining inhibition is then enriched in feedforward inhibition (FF-IPSC, Figure S4), although some reduction of FF inhibition could also occur due to its smaller effect on mPP terminals (Macek et al. 1996). In this condition, whole-cell voltage-clamp experiments were performed in mat-GCs upon stimulation of mPP, before and after optogenetic release of ACh (Figure S4G-H). Cholinergic modulation strongly reduced the FF-enriched-IPSC (Figure S4H-I), and the time course of its reduction parallels the potentiation of the population spike, and the total IPSC (Figure S2E). Interestingly, FB-IPSC shows a slower kinetics than the FF-enriched-IPSC (Figure 4B and S4H), suggesting that indeed both recording configurations recruit different inhibitory interneurons.

To further dissect the spatial contribution of inhibition that is affected by ACh, we isolated perisomatic and dendritic inhibition by placing a stimulating electrode either in the granule cell layer (GCL) or in the molecular layer. Each electrode evoked, as expected for the different location of inhibitory synapses, IPSCs with different kinetics (Figure 4D-E). Interestingly, we found that direct perisomatic inhibition was strongly reduced by endogenous ACh release, while dendritic inhibition was less affected (Figure 4E) and thus the perisomatic/dendritic IPSC ratio was largely reduced (Figure 4F), suggesting a spatial reorganization of inhibition.

### Acetylcholine strongly reduces PVIs but not SOMIs elicited IPSCs in granule cells

The reduction of specific components of inhibition by ACh suggests that its effect might be mediated by a specific subpopulation of inhibitory interneurons that express nicotinic receptors (Frazier et al., 2003; Griguoli and Cherubini, 2012; Prince et al., 2016). As previously mentioned, synaptic inhibition is provided by two main interneuron types in the hippocampus, PVIs and SOMIs (Freund and Buzsaki, 1996; Rudy et al., 2011; Yuan et al., 2017). PVIs provide powerful feedforward and feedback inhibition to large populations of GCs in the perisomatic region (Sambandan et al., 2010), while SOMIs are classically studied as dendritic targeting interneurons that participate more strongly in feedback inhibition, although it has been recently shown that they can be divided in functionally distinct subpopulations of SOMIs (Yuan et al., 2017). Because PVIs are the main FF inhibitory population in the DG (Hosp et al., 2014; Savanthrapadian et al., 2014), we hypothesized that ACh might act differentially on these two populations of interneurons.

In order to specifically assess the role of ACh in PVI-to-GC inhibition, a Cre-inducible AAV vector carrying ChR2 (AAV-CAG-DIO-ChR2) was stereotaxically injected in the DG of PV-Cre animals. We obtained hippocampal slices from these animals, in which we optogenetically activated PVIs (Figure 5A). We delivered light trains of 10 pulses at 10 Hz (PW = 5ms), while recording the evoked IPSCs in mat-GCs using patch-clamp whole-cell recordings at the reversal potential of excitation (∼0mV). Consistent with the main role of PVIs in the feedforward inhibitory circuit of the DG, IPSCs induced on GCs by activation of PVIs were strongly reduced by bath application of CBC (Figure 5B and 5C). Interestingly, when we studied the effect of CBC onto SOMI-elicited-IPSCs (by injecting AAV-CAG-DIO-ChR2 in the DG of SOM-Cre animals, Figure 5D), the recorded IPSCs were not affected (Figure 5E and 5F), suggesting that ACh selectively reduces PVI-mediated inhibition onto GCs. Importantly, the IPSCs evoked by stimulation of PVIs have a faster kinetics than the ones evoked by stimulation of SOMIs, suggesting, as expected, that the synaptic location of inhibition mediated by PVI is closer to the soma than the inhibition mediated by SOMI (Figure 5B and 5E).

**Figure 5.**
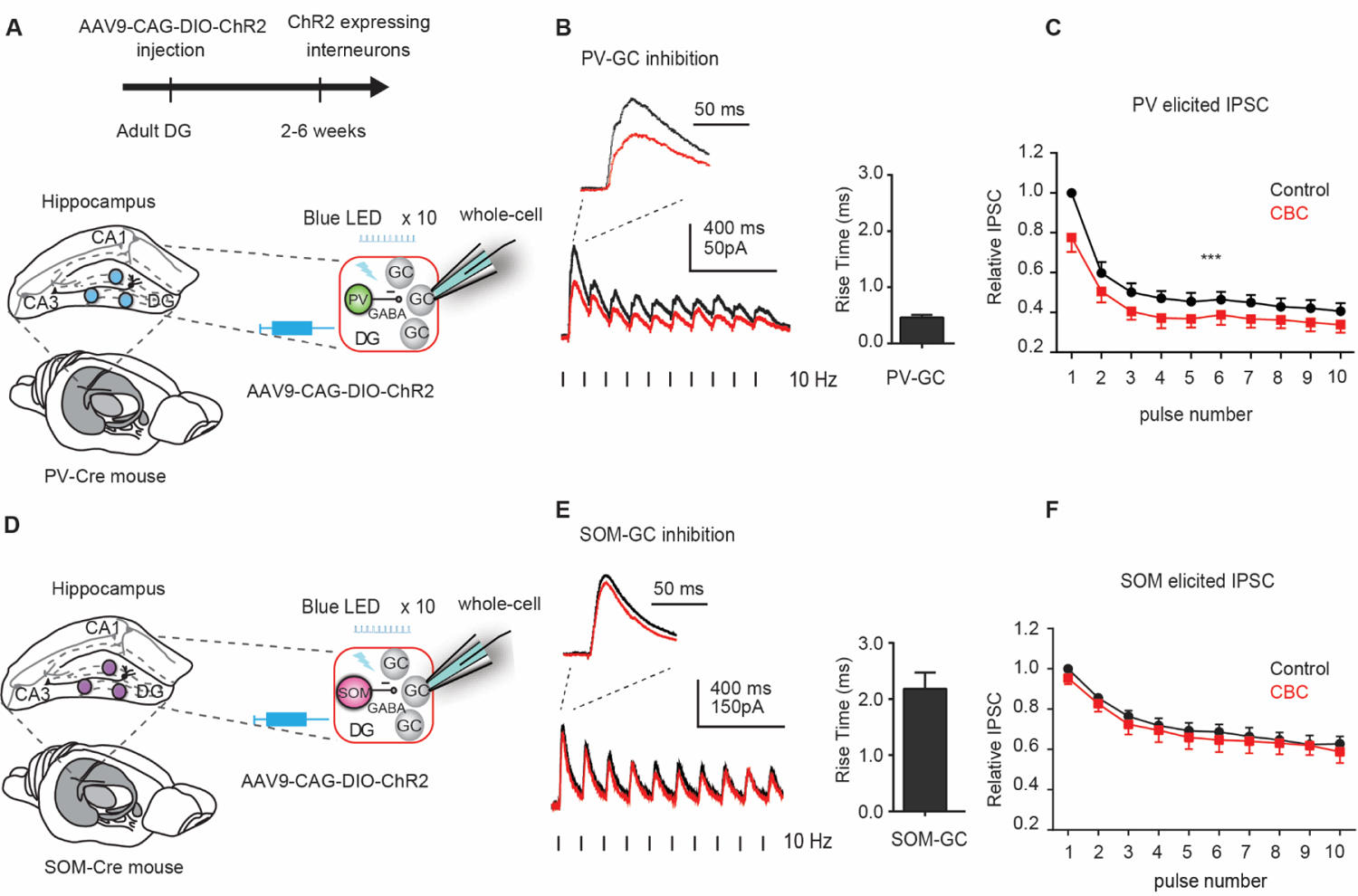
Acetylcholine reduces PVI but not SOMI elicited IPSCs in mat-GC. (A) Experimental scheme. PV-Cre mice were stereotaxically injected with the conditional ChR2 adenovirus (AAV-CAG-DIO-ChR2) in the DG to induce light evoked action potentials in PVIs from this region. The upper timeline indicates the time of viral injection. 2-6 weeks later, hippocampal slices were obtained from these animals. Blue light trains of 10 pulses at 10 Hz were delivered to the DG while whole-cell recordings were performed on mat-GCs at the reversal potential of excitation (0mV) to record light evoked IPSCs. (B) Example trace showing light evoked IPSCs in control condition (black) or after bath application of CBC (red). The dash indicates the time of the stimulus. Inset shows the same trace with increased scale. The graph on the right shows the 10-90% rise time of PV-evoked IPSC. (C) IPSC peak at each pulse relative to the 1st pulse in the control condition (black) or after bath application of CBC (red). CBC strongly reduced PV-to-GC IPSCs (two-way ANOVA, control vs CBC F _9, 45_ = 36.95: \*\*\**p* < 0.0001; variation in pulse number F _1, 5_ = 12.10: *** *p* = 0.0002; interaction F _9, 45_ = 1.356: *p* = 0.6227, N=9 cells from 3 mice). (D) Experimental scheme. The experimental design was identical to (A) but SOM-Cre animals were used in order to specifically activate this subpopulation of interneurons in the DG. (E) Example trace showing light evoked IPSCs in control condition (black) or after bath application of CBC (red). The dash indicates the time of the stimulus. Inset shows the same trace with increased scale. The graph on the right shows the 10-90% rise time of SOM-evoked IPSC. (F) IPSC peak at each pulse relative to the 1st pulse in the control condition (black) or after bath application of CBC (red). CBC did not change SOM-to-GC IPSCs (two-way ANOVA, control vs CBC F _1, 6_ = 1.403: *p* = 0.2810; variation in pulse number F _9, 54_ = 64.89: \*\*\**p* < 0.001; interaction F _9, 54_ = 0.6666: *p*= 0.7351, N=7 cells from 3 mice).

### Acetylcholine shifts the evoked E/I balance of PVIs and SOMIs in opposite directions

Because PVIs are recruited by mPP axons to produce feedforward inhibition, we further explored how that population of neurons processed afferent inputs upon cholinergic modulation. We performed whole-cell voltage-clamp recordings of PVIs to measure excitatory and inhibitory responses elicited at 10 Hz stimulation of the mPP (Figure 6A and 6B). Remarkably, in addition to the reduction of direct inhibition from PVIs to GCs by release of ACh, CBC produced a powerful decrease in the E/I balance of PVIs in response to mPP activation (Figure 6C and 6D). This results suggests that ACh can reduce PVI control of GC activity via at least two distinct but convergent mechanisms, an increased inhibition onto PVI (Figure 6A-D) and a reduction of PVI induced inhibition onto GCs (Figure 5A-C).

**Figure 6.**
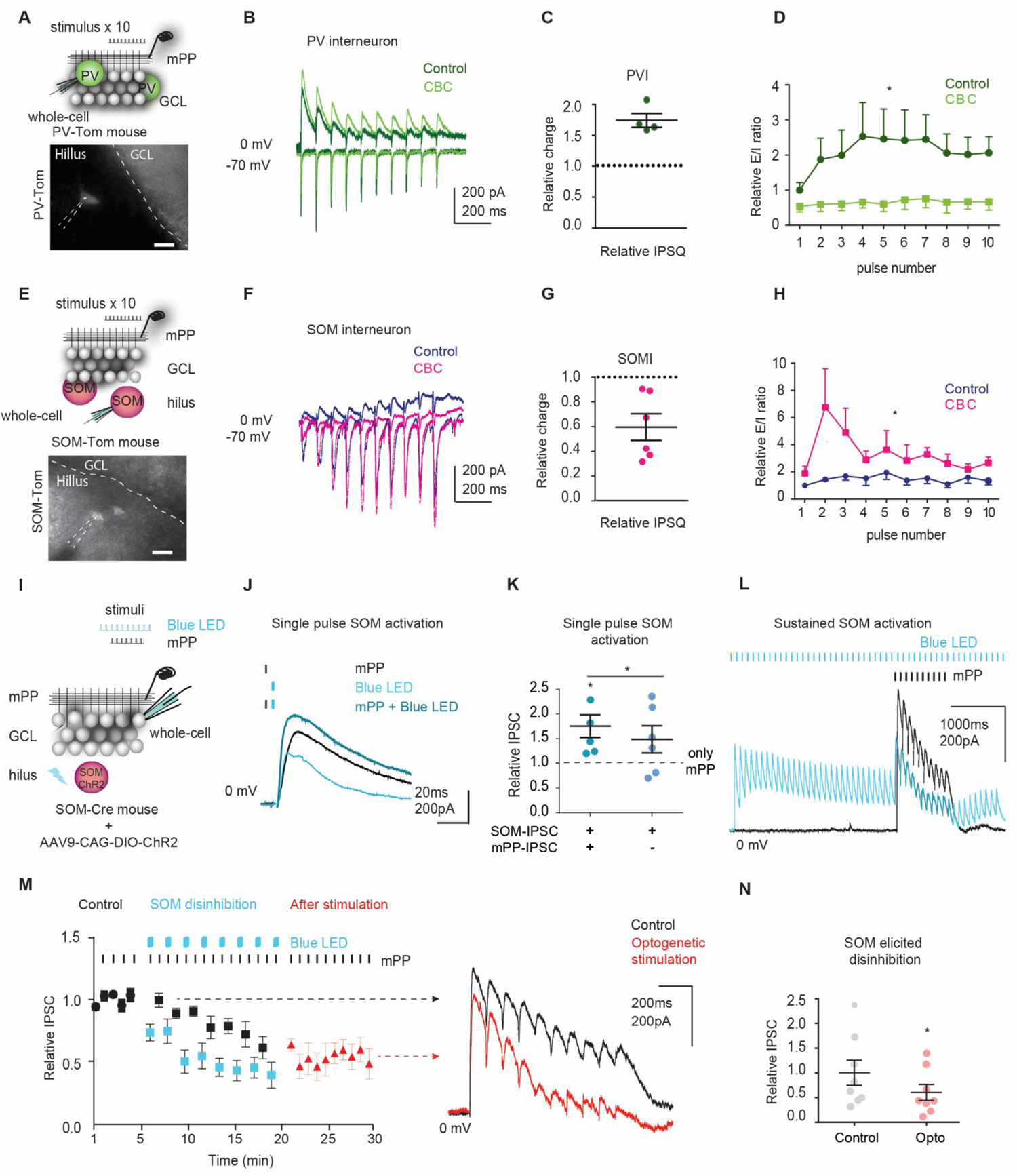
Cholinergic modulation recon figures PV and SOM inhibitory circuits. (A-H) *Cholinergic modulation shifts the evoked E/I balance of PVIs vs SOMIs in opposite directions*. (A) Recording configuration: Hippocampal slices were obtained from PV-Tom mice. Stimulation of mPP activates monosynaptic excitation and disynaptic inhibition onto PVIs. Whole-cell voltage-clamp recordings were obtained from PVIs in response to stimulation of mPP with trains at 10Hz. Example fluorescent image of a recorded PV interneuron in the hillus of a PV-Tom hippocampal slice. GCL, granule cell layer. Scale bar 20 um. (B) Example trace of evoked postsynaptic currents after mPP stimulation with a train of 10 pulses at 10Hz in control (dark blue) or after bath application of CBC (light blue). IPSCs were recorded at the reversal potential of excitation (0mV), and EPSCs were recorded at the reversal potential of inhibition (−70 mV). (C) CBC significantly increased the total evoked IPSQ in PVIs relative to control (Control vs CBC \*\**p*<0.01, one sample t-test, N=4 cells from 4 mice). (D) E/I balance at each pulse relative to the 1st pulse before (Control, dark blue) or after bath application of CBC (light blue). CBC significantly reduced the E/I balance of PV interneurons (two-way ANOVA, control vs CBC F _1, 30_ = 67.80: \*\*\**p*<0.001; variation in pulse number F _9, 30_ = 0.4328: *p* = 0.9065; interaction F _9. 30_ = 0.6301: *p*= 0.7622, N=4 cells from 4 mice). (E) Top, experimental scheme. Hippocampal slices were obtained from SOM-Tom mice. The recording configuration was the same as in (A). Example fluorescent image of a recorded SOM interneuron in the hillus of a SOM-Tom hippocampal slice. GCL, granule cell layer. Scale bar 20 um. (F) Example trace of evoked postsynaptic currents after mPP stimulation with a train of 10 pulses at 10Hz in control (purple) or after bath application of CBC (pink). IPSCs were recorded at the reversal potential of excitation (0mV), and EPSCs were recorded at the reversal potential of inhibition (−70mV). (G) CBC significantly decreased the total evoked IPSQ in SOMIs relative to control (Control vs CBC *p<0.05, one sample t-test, N=6 cells from 3 mice). (H) E/I balance at each pulse relative to the 1st pulse in control condition (purple) or after bath application of CBC (pink). CBC significantly increased the E/I balance of SOMIs (two-way ANOVA, control vs CBC F _1, 40_ = 24.32: \*\*\**p*<0.001; variation in pulse number F _9, 40_ = 1.227: *p* = 0.3066; interaction F _9, 40_ = 1.263: *p* = 0.2867, N=6 cells from 3 mice). (I-N) *SOM induced disinhibition of GCs*. (I) Experimental scheme. SOM-Cre mice were stereotaxically injected with the conditional ChR2 adenovirus (AAV-CAG-DIO-ChR2) in the DG to induce light evoked action potentials in SOMIs from this region. Hippocampal slices were obtained from these animals. Whole-cell recordings were performed on mat-GCs at the reversal potential of excitation (0mV) to record light and/or mPP evoked IPSCs. (J) Example IPSC traces obtained upon a single pulse stimulation of mPP (Black), SOMIs (Light blue) or both (Turquoise). (K) Mean peak IPSC values relative to activation of mPP only. Simultaneous co-activation of mPP and SOMIs produces sublinear summation of the IPSCs (one-way ANOVA, F _1, 5.2_ = 6.64: \**p*<0.05; post hoc Tukey comparisons: mPP vs mPP + SOMIs *p<0.05, N=6 cells from 3 mice). (L) Blue light trains of 5 seconds at 10 Hz were delivered to the DG to activate SOMIs, indicated by the blue dashed lines. mPP axons where electrically stimulated during SOM activation, indicated by the black dashed lines. Example traces of the IPSCs obtained during mPP stimulation (Black) or during both blue light and electrical stimulation (Turquoise). (M) Temporal evolution of the peak IPSC relative to control before (Black), during (Light Blue) and after (Red) optogenetic stimulation of SOMIs. The blue dashed line indicates the time of the sustained (5 seconds at 10Hz) activation of SOMIs. Right, example traces of the IPSCs obtained upon mPP stimulation in control (black) or after blue light stimulation protocol to activate SOMIs (Opto, red). The dashed line shows the time of each stimulus. (N) Activation of SOMIs strongly reduced mPP evoked IPSCs on mat-GCs (Control vs light stimulated: \**p*<0.05, paired t-test, N=8 cells from 3 mice).

The stronger inhibition received by PVIs (Figure 6C) upon cholinergic modulation suggests that another population of interneurons should be more active in this modulatory state. Because a subpopulation of SOMIs are known to inhibit PVIs (Yuan et al., 2017), we then performed whole-cell voltage-clamp recordings of hilar SOMIs to measure excitatory and inhibitory responses elicited by stimulation of the mPP (Figure 6E and 6F). Strikingly, CBC produced an increase in the E/I balance of SOMIs (Figure 6G and 6H), suggesting that these interneurons are more active during a cholinergic neuromodulatory state and could be mediating a disinhibitory effect onto the population of GCs.

These results, where ACh have opposite effects on SOMIs compared to PVIs, suggests that cholinergic modulation reconfigures inhibitory microcircuits shifting the relative contribution of different populations of interneurons that in turn control the DG activity.

### SOMIs activation can mimic cholinergic disinhibition

To evaluate the effect that the increased activity of SOMIs could have in the response of GCs to mPP stimulation, we optogenetically activated SOMIs while recording inhibitory currents on GCs in response to mPP stimulation (Figure 6I). Sustained activation of SOMIs previous to mPP stimulation reduced mPP evoked inhibition in GCs (Figure 6L-M), indicating that these interneurons are in fact able to produce disinhibition of GCs. Interestingly, activating SOMIs with a single light pulse at the same time of mPP stimulation does not reduce inhibition but in fact summates inhibition (Figure 6J-K), indicating the importance of a sustained activity to observe the disinhibitory effect. Remarkably, pairing repetitive stimulation of SOM with mPP several times produces a reduced IPSC that follows the same time course as ACh-mediated IPSC reduction and also outlasts SOM stimulation (Figure 6M-N). This suggests that concurrent mPP and SOMIs activation can induce a plasticity in inhibitory circuits that mimics cholinergic induced changes. Although SOMI-induced IPSC reduction was significant across all 10 pulses of the train, the effect was stronger for the last pulse compared to the first, which could give rise to a selective increase in gain at high frequency inputs (% of remaining IPSC for the first pulse: 74.9±20.0, last pulse: 31.1± 13.3, N =5 cells from 3 mice). These results show that activating SOMIs in the DG can bypass the necessity for cholinergic release while mirroring its effects.

### High-frequency stimulation produces LTP in the DG when paired with activation of cholinergic terminals

What is the functional consequence of the observed reconfiguration of inhibition? One possibility is that by disinhibiting GCs, ACh could produce a temporal window to allow plasticity in an otherwise very sparsely active region. We tested this hypothesis by using a tetanic stimulation protocol on hippocampal slices (Figure 7A). Importantly, this stimulation protocol is insufficient to produce LTP on mat-GCs unless inhibitory circuits are blocked (Snyder et al., 2001). We recorded the slope of the field excitatory postsynaptic potential (fEPSP) in slices obtained from ChAT-ChR2 mice, before and after tetanic stimulation (Figure 7B). As previously reported, this high frequency stimulation (HFS) was insufficient to produce a potentiation of the response, consistent with the fact that the vast majority of the DG GCs are mature. Remarkably, in the same slices we were able to induce LTP if we paired tetanic stimulation with endogenous release of ACh, which was not observed in Cre^-/-^ animals (Figure 7C). These results indicate that ACh can lower LTP-induction thresholds in the DG, providing a temporal window in which incoming inputs are able to induce long term synaptic changes.

**Figure 7.**
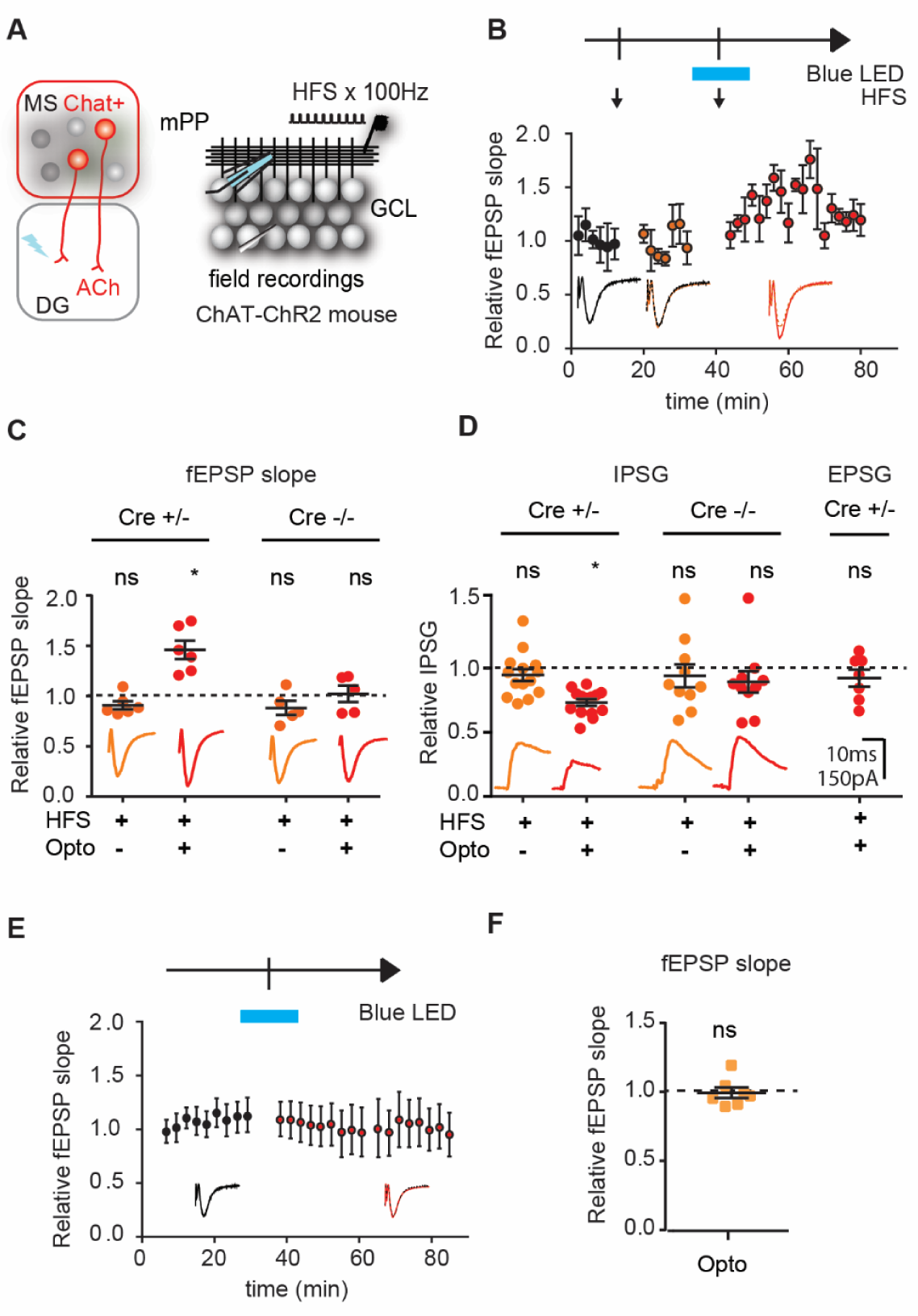
High frequency stimulation produces LTP in the DG when paired with optogenetic activation of cholinergic terminals. (A) Experimental scheme. Hippocampal slices were obtained from ChAT-ChR2 mice. A stimulating electrode was placed in the medial perforant path (mPP) to deliver a baseline (0.05 Hz) or a high frequency stimulation protocol (HFS, 4 trains, 500 ms each, 100 Hz within the train, repeated every 20 s) with or without blue light activation of cholinergic terminals. (B) Temporal evolution of the relative fEPSP slope along the experiment (average from 5 experiments). The black arrow indicates the time at which the HFS protocol was delivered; blue bar indicates the time at which blue light (4 trains, 5000 ms each, 100 Hz within the train repeated every 20s) was delivered. (C) Average fEPSP slope 10 min after HFS (HFS) and 10 min after pairing light and electrical stimulation (HFS + Light) relative to baseline, for Cre^+/-^ and Cre^-/-^ mice. HFS was insufficient to produce potentiation of the fEPSP unless paired with light activation of cholinergic axons. No potentiation was observed in Cre^-/-^ mice (one-way ANOVA F _3, 23_ = 13.99: \**p*<0.05, Bonferroni’s multiple comparisons: Baseline vs HFS Cre^+/-^ or Cre^-/-^ p>0.05; Baseline vs HFS + Light Cre^+/-^ \*\*\**p*<0.001; Baseline vs HFS + Light Cre^-/-^ p>0.05, N≥5 slices from 6 mice). (D) Same as in (C) but for excitatory and inhibitory conducances. Average peak IPSG or EPSG 10 min after HFS (HFS) and 10 min after pairing light and electrical stimulation (HFS + Light) relative to baseline, for Cre^+/-^ and Cre^-/-^ mice. Pairing HFS with activation of cholinergic fibers produced a reduction in the evoked IPSG in Cre^+/-^ mice, while EPSG remained unchanged. No change in conducances was observed in Cre^-/-^ mice (IPSG: One-way ANOVA F _3, 44_ = 3.15: \**p*<0.05, Tukey’s multiple comparisons: Baseline vs HFS Cre^+/-^ or Cre^-/-^ p>0.05; Baseline vs HFS + Light Cre^+/-^ \**p*<0.05; Baseline vs HFS + Light Cre^-/-^ p>0.05, N≥10 cells from 10 mice. EPSG Paired t-test: Baseline vs HFS + Light Cre^+/-^ *p*>0.05, N = 7 cells from 3 mice). (E) Temporal evolution of the relative fEPSP slope along the experiment (average from 6 slices). Electrical stimulation was kept constant at a frequency of 0,05Hz. Blue bar indicates the time at which blue light (4 trains, 5000 ms each, 100 Hz within the train repeated every 20s) was delivered. (F) Average fEPSP slope before (baseline), 10 min after pairing optogenetic stimulation. Optogenetic activation of cholinergic terminals was insufficient to produce potentiation of the fEPSP when the mPP was stimulated at low frequency (Paired t-test: Baseline vs Opto *p*>0.05, N=7 slices from 4 mice).

To evaluate if the high frequency protocol that produced LTP when paired with ACh release also produced a decrease in inhibition, we recorded IPSCs and EPSCs during the LTP experiment (Figure 7D). We observed that the HFS produced a reduction of inhibition when paired with endogenous release of ACh, which was not observed in Cre ^-/-^ animals, while excitation remained unchanged (Figure 7D). Remarkably, the same amount of activation of cholinergic terminals does not induce LTP if low-frequency stimuli are delivered to the mPP, indicating that ACh could act as a selective amplifier for high frequency inputs (Figure 7E and 7F).

## DISCUSSION

In the present work, we showed that endogenous ACh release into the DG produced an increased activation of mature GCs in response to afferent stimuli, without changing the activation of immature GCs. The increased response is due to a highly precise reconfiguration of inhibitory circuits that results in a net disinhibition of mature GCs, creating a window of opportunity for plasticity. In accordance with this, coincident activation of afferent perforant path inputs with ACh release produced a long-term-potentiated response in GCs.

The difference in the cholinergic effect on mature and immature GCs can be explained by the fact that immature neurons have underdeveloped perisomatic feedforward inhibition (Marín-Burgin et al., 2012), which is presumably the main target of the cholinergic effects. The cholinergic system is then able to independently modulate the input processing properties of the mat-GC population, without interfering with the parallel processing of the sub-population of immature neurons.

Our results show a highly specific reconfiguration of inhibitory microcircuits, producing opposite effects in different subpopulations of genetically defined interneurons. We observed a nicotinic-dependent reduction of inhibition in mat-GCs. In the DG, feedforward inhibition is primarily carried out by PVIs (Sambandan et al., 2010; Zipp et al., 1989). PVIs received more mPP evoked inhibition under cholinergic modulation but also produced less direct inhibition onto mat-GCs. These two distinct mechanisms result in a reduction of PV mediated control of GC activity.

Cholinergic reduction of inhibition has also been observed in CA1, where inhibition from PVIs to pyramidal neurons is reduced by ACh through nicotinic receptors (Tang et al., 2011), suggesting a common mechanism of cholinergic-mediated disinhibition in the hippocampus. Interestingly, despite being nicotinic dependent, cholinergic disinhibition of GCs and the corresponding increase in the population spiking activity both ramp up slowly, a phenomenon observed in other studies that found slow long-lasting effects mediated by mechanisms that include nicotinic receptors (Ballinger et al., 2016; Chen et al., 2015; Cheng and Yakel, 2015; Jiang et al., 2016).

Interestingly, a strong nicotinic mediated depolarization has been observed in hilar interneurons whose dendritic trees are within the hilus (Pabst et al., 2016). Hilar interneurons, including one subgroup of SOMIs, are known to inhibit PVI as well as other SOMIs (Yuan et al., 2017), being able to mediate the observed disinhibitory effect. Upon cholinergic modulation, we found an increase in the E/I balance of hillar SOMIs in response to afferent stimulation, which supports these neurons as good candidates for mediating the disinhibitory effect on mat-GCs. In fact, we observed that optogenetic stimulation of SOMIs reduces inhibition onto GCs in response to mPP stimulation, indicating that these interneurons do produce disinhibition in the DG, as has been also observed in CA1 (Leao et al., 2012). In summary, we propose that ACh reduces the influence of PVI onto GCs by two convergent actions: reducing the inhibition from PVI onto GC and, in addition, activating SOM neurons that inhibit PVI, further disinhibiting GC (Figure S5). Although our results can be explained by the antagonistic effects of ACh onto PVIs and SOMIs, the connectivity of different interneuronal subpopulations in the DG -as well as other brain regions-is emerging as more complex and diverse than usually described (Yuan et al 2017). Thus, other types of dentate interneurons or subgroups within these populations could also be affected by cholinergic modulation that need to be evaluated in future experiments.

The synaptic location of inhibition has important consequences for neuronal computations, where somatic inhibition modulates the gain (division effect) and dendritic inhibition have a subtractive effect (Pouille et al., 2013). The independent modulation of perisomatic and dendritic inhibition could presumably produce very different outputs of neuronal processing. Interestingly, *in vivo* studies have shown that inhibition at GC somata (presumably by PVIs) is reduced during exploratory movement, when ACh levels are high, whereas inhibition in the dendrites might be increased (Paulsen and Moser, 1998); These apparently contradictory changes could together act as filter and amplifier, increasing the contrast between signals with different relations to ongoing behavior (Paulsen and Moser, 1998). In this scenario, weak inputs would not be able to bypass the strong dendritic inhibition, but once they do, they are amplified by a reduced perisomatic inhibition. Our results are compatible with this model, in which we observed that the relative contribution of PVI-mediated perisomatic inhibition is reduced in comparison with dendritic inhibition, very likely carried by SOMIs. In fact, a number of studies propose a main role for ACh in increasing the signal to noise ratio (Hasselmo, 2006).The architecture of inhibitory subcircuits ensures richness in the possible dynamics within networks of principal neurons, and dictates the formation and maintenance of neuronal ensembles (Hu et al., 2014; Isaacson and Scanziani, 2011; Kepecs and Fishell, 2014). Through neuromodulation, different populations of interneurons can serve to gate information flow within the DG in specific behavioral events.It has been reported that LTP is more easily induced in the DG during exploration of novel environments, where ACh levels are high (Davis et al., 2004). Furthermore, Fos expression in the DG is greater after novel context exploration (VanElzakker et al., 2008). In the present work, we showed that pairing brief optogenetic activation of cholinergic fibers with stimulation of afferent inputs was sufficient to produce a strong potentiation of the mPP-GC pathway.

Because this potentiation requires more than one pulse of afferent stimulation, the cholinergic-DG system might serve to filter out temporally sparse inputs, conveying specificity and at the same time allowing high frequency inputs to produce long-term synaptic changes, thus increasing the signal to noise ratio of the circuit. That way a basal sparse code can be maintained while at the same time synapses receiving stronger stimuli together with ACh can be potentiated, acting as coincident detector.

This phenomenon is compatible with the phasic acetylcholine release in the hippocampus found during performance on a working memory task, where it is associated with the reward delivery, novelty, or fear (Acquas et al., 1996; Marrosu et al., 1995; Teles-Grilo Ruivo et al., 2017). In addition to pharmacological studies showing that nicotine infusions into the dorsal hippocampus produce dose-dependent enhancement of contextual fear conditioning (Davis et al., 2007; Kutlu and Gould, 2016), recent experiments using optogenetic release of ACh in the DG have shown an increase in associative fear memory in animals preexposed to a spatial context in the presence of ACh, suggesting that novelty-induced ACh release primes future contextual associations (Hersman et al., 2017).

The present work provides a circuit mechanism by which ACh dynamically reconfigures the DG network, allowing this brain region to change its information processing properties by shifting the relative contribution of different subpopulations of interneurons (Figure S5). These changes increase the output of DG and also favor the induction of long-term synaptic plasticity, essential for memory formation. This modulation can serve *in vivo* to adapt the encoding to the behavioral and cognitive demands of different tasks. Understanding the role of cholinergic modulation of input processing in the hippocampus can have important implications for neurodegenerative diseases in which the cholinergic and consequently the memory systems are severely impaired as observed in Alzheimeŕs disease (Ballinger et al., 2016).

## ACKNOWLEDGEMENTS

We thank members of the Marin-Burgin lab, the Refojo lab and Muraro lab for insightful discussions. We thank Jens Bruening for sharing Ai27 mice with us. We thank Hillel Adesnik for his valuable comments on the manuscript. This work was supported by grants from the Argentine Agency for the Promotion of Science and Technology (PICT2013-0182 and PICT2015-0364, PICT2018-0880), ECOS 2014, IDRC108878 to AMB and FOCEM-Mercosur (COF 03/11).

## AUTHOR CONTRIBUTION

MBO performed most experiments, analysis and contributed with the writing of the manuscript and experimental design. OP performed part of the electrophysiological experiments and analysis. DR, NF and SAR contributed with electrophysiological analysis and manuscript writing. GL and LAB contributed with the histochemistry of ChAT mice. AMB designed the research and writing of the manuscript.

## STAR METHODS

### KEY RESOURCES TABLE

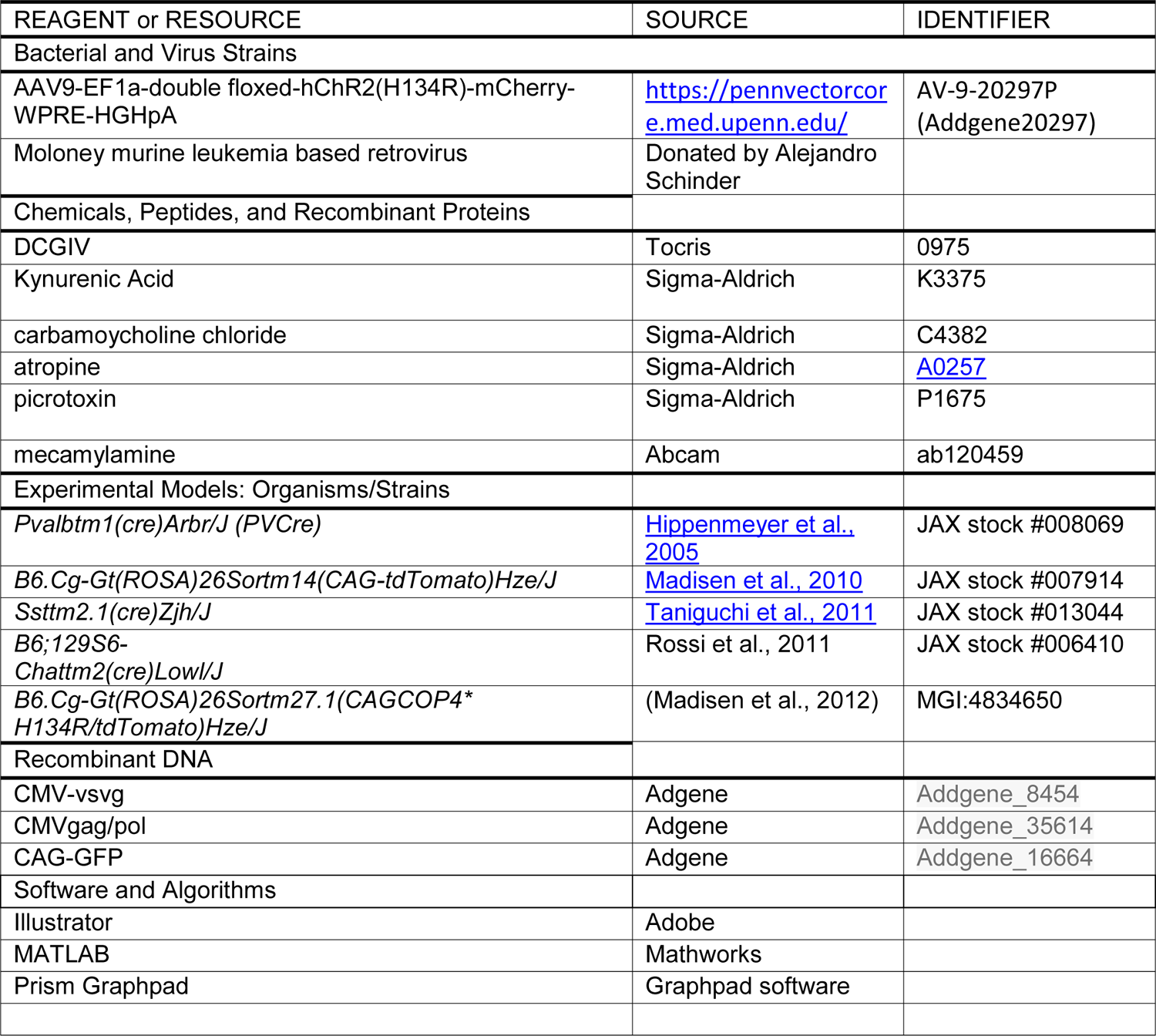

### LEAD CONTACT AND MATERIALS AVAILABILITY

Further information and requests for resources and reagents should be directed to and will be fulfilled by the Lead Contacts, Antonia Marin-Burgin and Mora Ogando. This study did not generate new unique reagents.

### EXPERIMENTAL MODEL AND SUBJECT DETAILS

#### Animals used

To generate *Pvalb^Cre^*;CAG^FloxStopTom^ (PV-Tom) mice, *B6;129P2-Pvalbtm1(cre)Arbr/J (PV^Cre^)* mice (Hippenmeyer et al., 2005) were crossed to *B6.Cg-Gt(ROSA)26Sortm14(CAG-tdTomato)Hze/J (Ai14)* conditional reporter mice. Animals heterozygous for Cre were used.

To generate *SOM^Cre^*;CAG^FloxStopTom^ (SOM-Tom) mice, *Ssttm2.1(cre)Zjh/J (SOM^Cre^)* were crossed to *B6.Cg-Gt(ROSA)26Sortm14(CAG-tdTomato)Hze/J (Ai14)* conditional reporter mice. Animals heterozygous for Cre were used.

To generate *ChAT^Cre^*;Ai27 (ChAT-ChR2) mice, *B6;129S6-Chattm2(cre)Lowl/J(ChAT^Cre^)* mice (Dannenberg et al., 2019; Herman et al., 2016; Yaeger et al., 2019) were crossed to *B6.Cg-Gt(ROSA)26Sortm27.1(CAG-COP4*H134R/tdTomato)Hze/J (Ai27)* conditional channelrhodopsin mice (Madisen et al., 2012). Animals heterozygous for Cre were used.

Experimental protocol (2020-03-NE) was evaluated by the Institutional Animal Care and Use Committee of the IBioBA-CONICET according to the Principles for Biomedical Research involving animals of the Council for International Organizations for Medical Sciences and provisions stated in the Guide for the Care and Use of Laboratory Animals. Mice were housed under controlled environment in a 12h light/dark cycle, with food and water ad libitum.

## METHOD DETAILS

### Animals and surgery for viral delivery

For the neurogenesis experiments, female C57Bl/6J mice 6–8 weeks of age were housed at 4 mice per cage, with one running wheel to increase neurogenesis. Running wheel housing started 2–4 days before surgery and continued until the day of slice preparation to increase neurogenesis. Although the running wheel may cause some differences in the maturation stages (Piatti et al., 2011), those are mostly in the ventral hippocampus that was not studied in the experiments. For surgery, mice were anesthetized (150 µg ketamine/15 µg xylazine in 10 µl saline/g), and virus (1 µl at 0.125 µl/min) was infused into the dorsal area of the right DG using sterile microcapillary calibrated pipettes and stereotaxic references (coordinates from bregma: −2 mmanteroposterior,-1.5 mm lateral, −1.9 mm ventral). Animals were killed for acute slice preparation 4 weeks after the surgery. While all experiments involving 4wpi-GCs were performed with mice under the running wheel housing condition, no differences were observed in the cholinergic modulation effects encountered in this study for the mat-GC population by this housing condition. We cannot, however, discard a possible modulation of septo-hippocampal circuits by running wheel housing.

For optogenetic experiments with interneurons, PV^Cre^ or SOM^Cre^ mice 8–20 weeks of age were anesthetized (150 µg ketamine/15 µg xylazine in 10 µl saline/g), and AAV9-EF1a-double floxed-hChR2(H134R)-mCherry-WPRE-HGHpA (AAV-DIO-ChR2) virus (0.5 µl at 0.125 µl/min) was infused into the dorsal area of the right DG using sterile microcapillary calibrated pipettes and stereotaxic references (same coordinates as before). Animals were killed for acute slice preparation 2-6 weeks after the surgery.

For optogenetic activation of septal cholinergic fibers in the DG, ChAT^Cre^ mice 8-16 weeks of age were anesthetized (150 µg ketamine/15 µg xylazine in 10 µl saline/g), and AAV-DIO-ChR2 virus (1 µl at 0.125 µl/min) was infused into the medial septum using sterile microcapillary calibrated pipettes and stereotaxic references (coordinates from bregma: 0.9 mm anteroposterior, 0.0 mm lateral, −3.6 mm ventral). Animals were killed for acute slice preparation 2-6 weeks after the surgery.

### Viral vectors preparation

A replication-deficient retroviral vector based on the Moloney murine leukemia virus was used to express GFP under a CAG promoter as previously described (Marín-Burgin et al., 2012). Retroviral particles were assembled using three separate plasmids containing the capside (CMV-vsvg), viral proteins (CMV-gag/pol), and the transgene (CAG-RFP or CAG-GFP). Plasmids were transfected onto HEK 293T cells using deacylated polyethylenimine. Virus-containing supernatant was harvested 48 hr after transfection and concentrated by two rounds of ultracentrifugation. Virus titer was typically ∼10^5^ particles/μl.

### Slice preparation

Mice were anesthetized and decapitated. Brains were removed into a chilled solution containing (mM) 110 choline-Cl^−^, 2.5 KCl, 2.0 NaH2PO_4_, 25 NaHCO_3_, 0.5 CaCl_2_, 7 MgCl_2_, 20 dextrose, 1.3 Na^+^-ascorbate, 3.1 Na^+^-pyruvate. The right hippocampus was dissected and slices (400 µm thick) were cut transversally to the longitudinal axis in a vibratome and transferred to a chamber containing artificial cerebrospinal fluid (ACSF; mM): 125 NaCl, 2.5 KCl, 2.3 NaH_2_PO_4_, 25 NaHCO_3_, 2 CaCl_2_, 1.3 MgCl_2_, 1.3 Na^+^-ascorbate, 3.1 Na^+^-pyruvate, and 10 dextrose (315 mOsm). Slices were bubbled with 95% O_2_/5% CO_2_ and maintained at 30°C for approximately 1 hr before experiments started. Salts were acquired from Sigma-Aldrich (St. Louis, MO). When carbamoycholine chloride (CBC, 5-20 µM, Sigma Aldrich), atropine (10 µM, Sigma Aldrich), mecamylamine (20 µM, ABCAM), picrotoxine (PTX, 100 µM, Sigma Aldrich), were used, they were perfused into the bath solution.

### Extracellular recordings

A field-recording microelectrode was placed on the granule cell layer (GCL) or molecular layer (ML) to record the population spike or the fEPSP, respectively, in response to the mPP stimulation. The input strength was adjusted to elicit population spike responses between 0.5 and 2 mV or fEPSP responses between −0.15 and −0.6 mV/ms, within the linear part of the dynamic range of responses. For a given input strength within a slice, the number of mPP axons stimulated in control conditions or after neuromodulatory manipulations were the same.

### Electrophysiological single cell recordings

Recorded neurons were visually identified by fluorescence and infrared DIC video microscopy. The mature neuronal population encompassed RFP^−^ or GFP^−^ neurons localized in the outer third of the GCL (Mongiat et al., 2009). Whole-cell recordings were performed using microelectrodes (4–5 MΩ) filled with (in mM) 130 CsOH, 130 D-gluconic acid, 2 MgCl_2_, 0.2 EGTA, 5 NaCl, 10 HEPES, 4 ATP-tris, 0.3 GTP-tris, 10 phosphocreatine. In experiments where the intrinsic responses to current pulses were evaluated, a potassium gluconate internal solution was used (in mM): 120 potassium gluconate, 4 MgCl_2_, 10 HEPES buffer, 0.1 EGTA, 5 NaCl, 20 KCl, 4 ATP-tris, 0.3 GTP-tris, and 10 phosphocreatine (pH = 7.3; 290 mOsm). In experiments where the perisomatic and dendritic inhibitory responses were evaluated, a CsOH with symmetric cloride internal solution was used (in mM): 140 CsCl, 2 MgCl2, 0.1 EGTA, 5 NaCl, 10 HEPES, 4 ATP-tris, 0.3 GTP-tris, 10 phosphocreatine (pH = 7.3; 290 mOsm). Loose-patch recordings were performed with ACSF-filled patch pipettes (5–6 MΩ). Field recordings were performed using patch pipettes (5 MΩ) filled with 3 M NaCl. Recordings were obtained using Multiclamp 700B amplifiers, (Molecular Devices, Sunnyvale, CA), digitized, and acquired at 20 KHz onto a personal computer using the pClamp10 software. Membrane capacitance and input resistance were obtained from current traces evoked by a hyperpolarizing step of 10 mV. Series resistance was typically 10–20 MΩ, and experiments were discarded if higher than 40 MΩ.

### Evoked postsynaptic currents and conductance

Evoked monosynaptic excitatory postsynaptic currents (EPSC) and inhibitory postsynaptic currents (IPSC) were recorded after stimulation. EPSCs were isolated by voltage clamping GC, PVI or SOMI at the reversal potential of the IPSC measured for each individual neuron (∼-60 mV). In turn, IPSCs were recorded at the reversal potential of the EPSC (∼0 mV). Both, IPSCs and EPSCs were recorded interleaved from the same cell. Direct monosynaptic IPSC elicited in presence of kynurenic acid (KYN, Sigma Aldrich) was subtracted from the total IPSC in all shown traces except Figure S4. Synaptic excitatory and inhibitory conductances were computed as the EPSC or IPSC divided by the driving force at which the synaptic currents were recorded.

### Measurement of feedforward enriched IPSC and feedback IPSC

To dissect feedforward-enrichedcomponent of the inhibitory response, the stimulating electrode was placed on the mPP, while mossy fiber synapses were blocked with DCGIV, an agonist of group II metabotropic glutamate receptors (mGluR2/3) that reduces release probability in mossy fiber terminals (1 μM, Tocris)(Kamiya et al., 1996). This strongly abolishes the feedback component of the IPSC (Figure S4). Although DCG4 could also affect release from mPP axons (Macek et al., 1996), which could impair recruitment of feedforward interneurons, we have previously shown that recruitment of PV neurons, that participate in FF-IPSC, is not affected by application of DCG4 at the intensities of stimulation we use, that is above threshold for interneurons recruitment (Pardi et al., 2015). In this condition, the remaining feedforward-enriched current was measured, before and after the optogenetic stimulating protocol, always in presence of DCGIV. In order to isolate the feedback component of the inhibitory response, the stimulating electrode was placed directly on the GCL. As stated before, direct monosynaptic IPSC recorded in KYN was subtracted from the IPSC in all traces, except in Figure S4, in which traces of total IPSC, DCGIV and KYN are shown. Stimulation of GCL (Figure 4 A-C) or mPP (Figure 3 D) afferents evoked similar percentage of direct inhibition (26 ± 4 % vs 29 ± 5 % for GCL vs mPP evoked IPSC respectively).

### Characterization of direct perisomatic and dendritic inhibition

To dissect dendritic component of the inhibitory response, stimulating electrodes were placed in the granule cell layer (GCL) and the molecular layer (ML) to activate monosynaptic perisomatic or dendritic inhibition. Experiments were performed in the presence of 6 mM kynurenic acid (Kyn) to block glutamatergic transmission. IPSCs were subsequently measured in whole-cell recordings from individual cells. In this conditions, perisomatic IPSCs display a reversal potential of ∼0 mV due to the recording conditions under symmetrical Cl^-^, while dendritic IPSCs display a positive driving force at that potential.

### Optogenetics

Hippocampal slices containing ChR2 expressed either in cholinergic fibers or interneurons were obtained. ChR2 expressing neurons or axons were stimulated using a 470 nm LED source delivered through the epifluorescence pathway of the upright microscope (63X objective for whole-cell recordings, 20X for field recordings) commanded by the acquisition software. For cholinergic neuromodulatory experiments in slices obtained from ChAT-ChR2 mice, 5 to 10 trains of blue light (10Hz, pulse width 10 ms) were delivered for 5 seconds every minute (Figure S2). Within the light train, either a single pulse or a train of 10 pulses at 10 Hz was delivered to the stimulating electrode (pulse width 0.2ms). Cholinergic modulation slowly augmented the amplitude of population spikes until reaching a plateau at around 10 minutes (Figure S2). We then measured the responses before and 15 minutes after the beginning of the light stimulation protocol. Both the control and the neuromodulated experimental groups were measured in the absence of light and at the same intensity of mPP stimulation, in order to compare two identically stimulated conditions.

In other experiments, slices were obtained from ChAT^Cre^ mice injected with the adenoviral vector AAV-DIO-ChR2 in the medial septum. 2-6 weeks after surgery, hippocampal slices were obtained from these animals and septal cholinergic axons in the DG were stimulated with blue light. For these experiments, light trains of 50 seconds were delivered to the preparation. At the onset of the light train, a train of 10 pulses of electrical stimulation was delivered at 10 Hz. Both the control and the neuromodulated experimental groups were measured in the absence of light, in order to compare two identically stimulated conditions.

For optogenetic activation of interneuron populations, PV-Cre or SOM-Cre animals were injected with AAV-DIO-ChR2 in the dorsal DG in order to express ChR2 specifically in either subpopulation of interneurons. Light trains of 10 pulses at 10 Hz (pulse width 1-5 ms) were delivered every minute. The intensity of the light was adjusted in order to have a non-saturating evoked IPSC.

### QUANTIFICATION AND STATISTICAL ANALYSIS

Unless otherwise specified, data are presented as mean ± SEM. Normality was assessed using Shapiro-Wilk’s test, D’Agostino & Pearson omnibus test, or Kolmogórov-Smirnov’s test, at a *p* value of 0.05. For normal distributions, homoscedasticity was assessed using Bartlett’s test and F-test, at a *p* value of 0.05. Two-tailed ordinary, one sample or paired t-test was used for single comparisons, and ordinary or repeated measures ANOVA for multiple comparisons, with post-hoc Tukey’s test.

## DATA AND CODE AVAILABILITY

The published article includes all datasets/codes generated or analyzed during this study.

