## Supplementary material for "Cholinergic modulation of dentate gyrus processing through dynamic reconfiguration of inhibitory circuits"

**This PDF file includes**

**Figures S1-S5.**

**Supporting Online Material**

**Figure S1. Carbachol increases evoked spiking activity in CA3.**

(A) Recording configuration. Hippocampal slices were obtained from adult mice. The scheme shows the recording configuration: a stimulating electrode was placed in the medial perforant path (mPP) to deliver 10 stimuli at 10 Hz; mPP axons activate GCs in the DG that in turn activate CA3 pyramidal neurons. Extracellular recordings were obtained from the both the granule cell layer (GCL) and pyramidal layer of CA3 region to detect population spikes.

(B) Example traces showing the population spike response to a train of 10 pulses at 10 Hz in DG (the asterisk denotes the amplitude of the population spike at each pulse) before (Control, black) or after bath application of CBC (red).

(C) Mean population spike amplitude in response to the 10 Hz train in control condition (black) and after bath perfusion of CBC (red). CBC increased the population spike response of GCs (Paired t-test control vs CBC, ***$p<0.001$ N = 5).

(D) Example traces showing the population spike response to a train of 10 pulses at 10 Hz in the pyramidal layer (the asterisk denotes the amplitude of the population spike at each pulse) before (Control, black) or after bath application of CBC (red).

(E) Mean population spike amplitude in response to the 10 Hz train in control condition (black) and after bath perfusion of CBC (red). CBC increased the population spike response of pyramidal neurons (Paired t-test control vs CBC, ***$p<0.001$ N=5).

**Figure S2. Optogenetic illumination protocol to evoke endogenous release of acetylcholine and time course of cholinergic modulation**

(A) Optogenetic illumination protocol: trains of blue light (10Hz, pulse width 10 ms) were delivered for 5 seconds every minute, beginning at second -2 with respect to the onset of mPP electrical stimulation. A train of 10 pulses at 10 Hz was delivered to the stimulating electrode (pulse width 0.2ms) so that the onset of each blue light pulse was coincident with each pulse of the electrical

stimulation. Because the total duration of the 10 pulse mPP stimulation is 1 second, the light stimulation outlasted the electrical mPP stimulation by 2 seconds.
(B) Field recordings from the GCL at increasing stimulus intensities. The black trace corresponds to the intensity which evokes the maximum population spike (100% population spike). The dotted green line indicates the slope of the fEPSP corresponding to the maximum population spike, considered 100% fEPSPslope.
(C) Left, recording configuration: in slices obtained from ChAT-ChR2 mice, we used blue light to activate the cholinergic terminals in the DG. A stimulating electrode was placed in the medial perforant path (mPP) to deliver 10 stimuli at 10 Hz; Extracellular recordings were obtained from the GCL to detect population spikes. Right, example traces showing the population spike response to a train of 10 pulses at 10 Hz before (control, black traces), during (dark red traces under light blue bar) or after optogenetic stimulation protocol to activate cholinergic terminals (red).
(D) Mean population spike amplitude relative to baseline before (Black) during (Red) or after optogenetic stimulation protocol for ChAT-ChR2 (Blue, N≥6) or ChAT-Tom (Gray, N=5). Endogenous release of ACh slowly increased the PS amplitude until reaching a plateau at 10 min in ChAT-ChR2 but not in ChAT-Tom mice. Potentiation remained significant even 10 minutes after optogenetic stimulation (23 ± 7 % increase vs baseline, *$p<0.05$ paired t-test, N = 6).
(E) Total (Blue, N=4) or FF-enriched (Green, N=5) mean IPSC amplitude relative to baseline before, during or after optogenetic stimulation protocol. Endogenous release of ACh slowly decreased the total and FF-enriched IPSC amplitude.
(F) Total IPSC in the presence of mecamylamine (Black, N=4) or FB-IPSC (Gray, N = 6) mean amplitude relative to baseline before, during, and after illumination protocol. Endogenous release of ACh does not alter the evoked IPSC in these conditions.

**Figure S3. Endogenous release of septal Ach increases spiking response in the DG.**

(A) Experimental design. The upper timeline indicates the time of adenoviral (AAV-DIO-ChR2) stereotaxic injection into the medial septum of ChAT-Cre mice.

2-6 weeks after the surgery acute hippocampal slices were obtained from these animals for specific activation of septohippocampal cholinergic projections. Right, the scheme shows the recording configuration: a stimulating electrode was placed in the medial perforant path (mPP) to deliver 10 stimuli at 10 Hz; Extracellular recordings were obtained from the GCL to detect population spikes.
(B) Example traces showing the population spike response to a train of 10 pulses at 10 Hz before (control, upper black) or after optogenetic stimulation protocol (red) (the asterisk indicates the peak of the population spike, when present, at each pulse).
(C) Average amplitude of the population spike at each pulse of the 10 Hz train in control (black) and after optogenetic stimutation protocol (red). Optogenetic stimulation increased population spike responses in hippocampal slices obtained from ChAT-Cre$^{+/-}$ mice injected with AAV-DIO-ChR2 (two-way ANOVA, variation between control and optogenetic stimulated $F_{1, 40} = 30.15$: ***$p < 0.0001$; variation between pulses $F_{9, 40} = 27.47$: ***$p < 0.0001$; interaction $F_{9, 40} = 8.599$: **$p < 0.01$ N=5 slices from 3 mice).

**Figure S4. ACh release reduces feedforward-enriched IPSC in mat-GCs.**
(A) Experimental scheme. Hippocampal slices were obtained from WT mice. Recording configuration showing the activated circuit. Stimulation of mPP-activated monosynaptic excitation and feedforward inhibition (FFI). Feedback inhibition (FBI) is recruited by the activity of GCs. Application of DCGIV strongly reduced feedback IPSC leaving the feedforward-enriched IPSC.
(B) Example traces of the total (Purple), FF-enriched (Black) and putative FB (Gray) IPSCs. FB-IPSC can be calculated by subtracting the IPSC in the presence of DCGIV from the total IPSC.
(C) Rise time (10-90%) of the FF-enriched and the arithmetically calculated FB-IPSCs. FF-enriched IPSC was significantly faster than the remaining FB IPSC (paired t-test, *$p<0.05$, N= 4 cells from 3 mcie).
(D) Schematic diagram of the recording configuration. Stimulating electrodes were placed in the GCL and the ML to disynaptically activate FB-enriched or total (FB + FF) IPSC, respectively. DGCIV was bath applied to compare the degree of reduction of total (ML evoked) and FB-enriched (GCL evoked) IPSC. Importantly,

kynureic acid was applied at the end of each recording in order to subtract the direct monosynaptic IPSC.

(E) Example traces recorded in the same cell for ML (top) and GCL (bottom) evoked IPSC in control conditions (total IPSC, Purple), after bath perfusion of DCGIV (Black) and after bath perfusion of kynurenic acid (Gray). The remaining IPSC after bath application of kynurenic acid is direct monosynaptic inhibition that was subtracted from all the traces at the end of the recording.

(F) DGCIV preferentially reduces GCL evoked IPSC. (T-test, ** $p<0.01$, N≥7 cells from 6 mice).

(G) Experimental scheme. Hippocampal slices were obtained from ChAT-ChR2 mice. Blue light stimulates cholinergic terminals in the DG. Recording configuration showing the activated circuit. Stimulation of mPP-activated monosynaptic excitation and feedforward inhibition (FFI). Feedback inhibition (FBI) is recruited by the activity of GCs. Application of DCGIV strongly reduced feedback IPSC leaving the feedforward enriched IPSC (FFe-IPSC).

(H) Example trace of evoked FF-enriched-IPSC after mPP stimulation with a train of 10 pulses at 10 Hz in control (black), or after blue light stimulation protocol to activate cholinergic terminals (red). Right, mean value of the FF-IPSC rise time (10-90%) from the recorded neurons. Upper inset shows the evoked FF-IPSC in an increased scale.

(I) Total inhibitory charge (IPSQ) evoked by the train. Light stimulation significantly reduced the FFe-IPSQ (control vs light stimulated: ** $p<0.01$, paired t-test, N=5 cells from 2 mice).

**Figure S5. Cholinergic reconfiguration of DG microcircuit processing model.**

(A) Cartoon showing the anatomic location of the main neural populations studied within the DG. When inputs arrive from the EC through the mPP axons, PVI are strongly activated (both directly from the mPP axons and indirectly through GCs) and inhibit GCs. Because of this strong inhibition, the output activity pattern of the GCL is relatively small.

(B) Activation of the afferent pathway and simultaneous optogenetic release of endogenous acetylcholine produces disinhibition of GCs by two convergent mechanisms. 1. ACh decreases PV-GC inhibition (which is mainly perisomatic). 2. ACh increases activation of PV-targeting SOMI, further reducing their control of GC activity. As a result, the output from GCs is increased.

ML = Molecular Layer. GCL = Granule Cell Layer. mPP = medial Perforant Path axons.

Figure S1

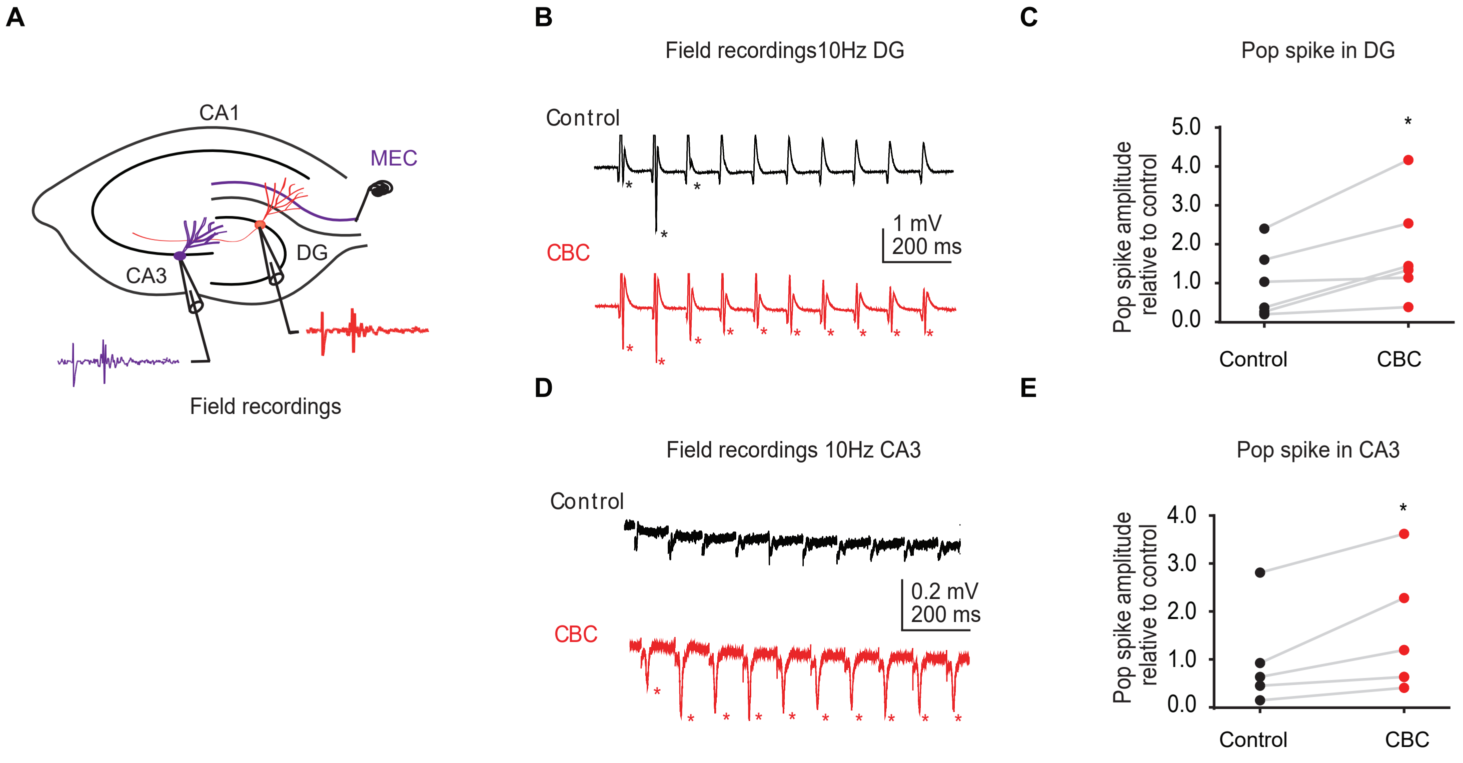
A
CA1
MEC
CA3
DG
Field recordings
B
Field recordings10Hz DG
Control
CBC
1 mV
200 ms
C
Pop spike in DG
Pop spike amplitude relative to control
5.0
4.0
3.0
2.0
1.0
0.0
Control
CBC
D
Field recordings 10Hz CA3
Control
CBC
0.2 mV
200 ms
E
Pop spike in CA3
Pop spike amplitude relative to control
4.0
3.0
2.0
1.0
0.0
Control
CBC

Figure S2

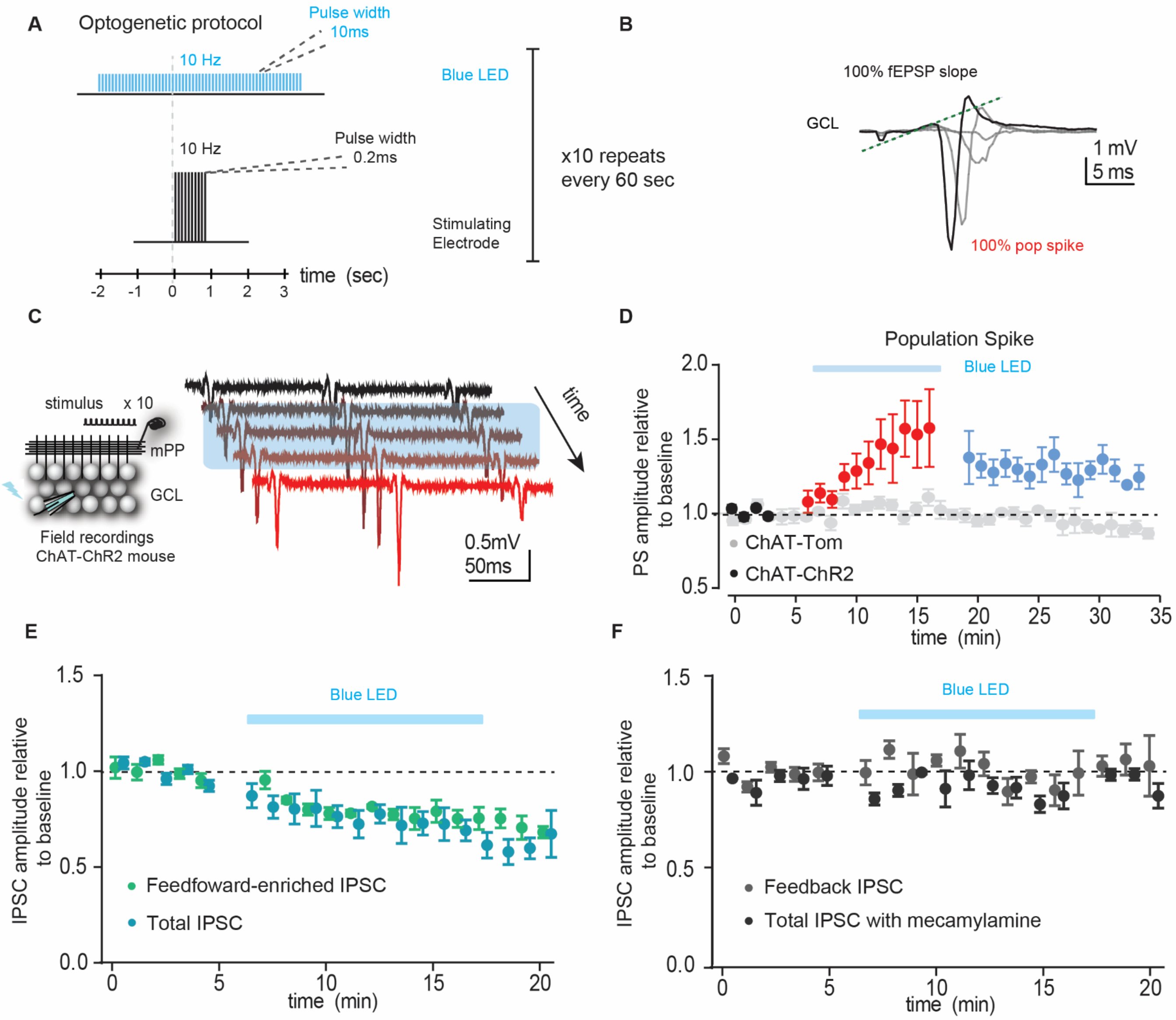

A
Optogenetic protocol
Pulse width
10ms
10 Hz
Blue LED
10 Hz
Pulse width
0.2ms
x10 repeats
every 60 sec
Stimulating
Electrode
time (sec)
-2 -1 0 1 2 3
B
100% fEPSP slope
GCL
1 mV
5 ms
100% pop spike
C
stimulus x 10
mPP
GCL
Field recordings
ChAT-ChR2 mouse
time
0.5mV
50ms
D
Population Spike
Blue LED
PS amplitude relative to baseline
ChAT-Tom
ChAT-ChR2
time (min)
E
Blue LED
IPSC amplitude relative to baseline
Feedfoward-enriched IPSC
Total IPSC
time (min)
F
Blue LED
IPSC amplitude relative to baseline
Feedback IPSC
Total IPSC with mecamylamine
time (min)

Figure S3

**A**

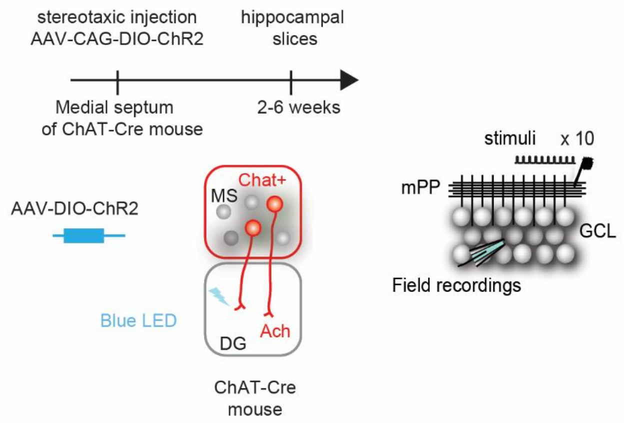


**B**

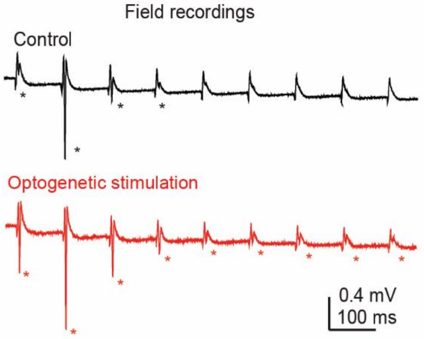


**C**

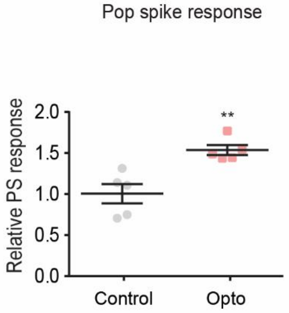

Figure S4

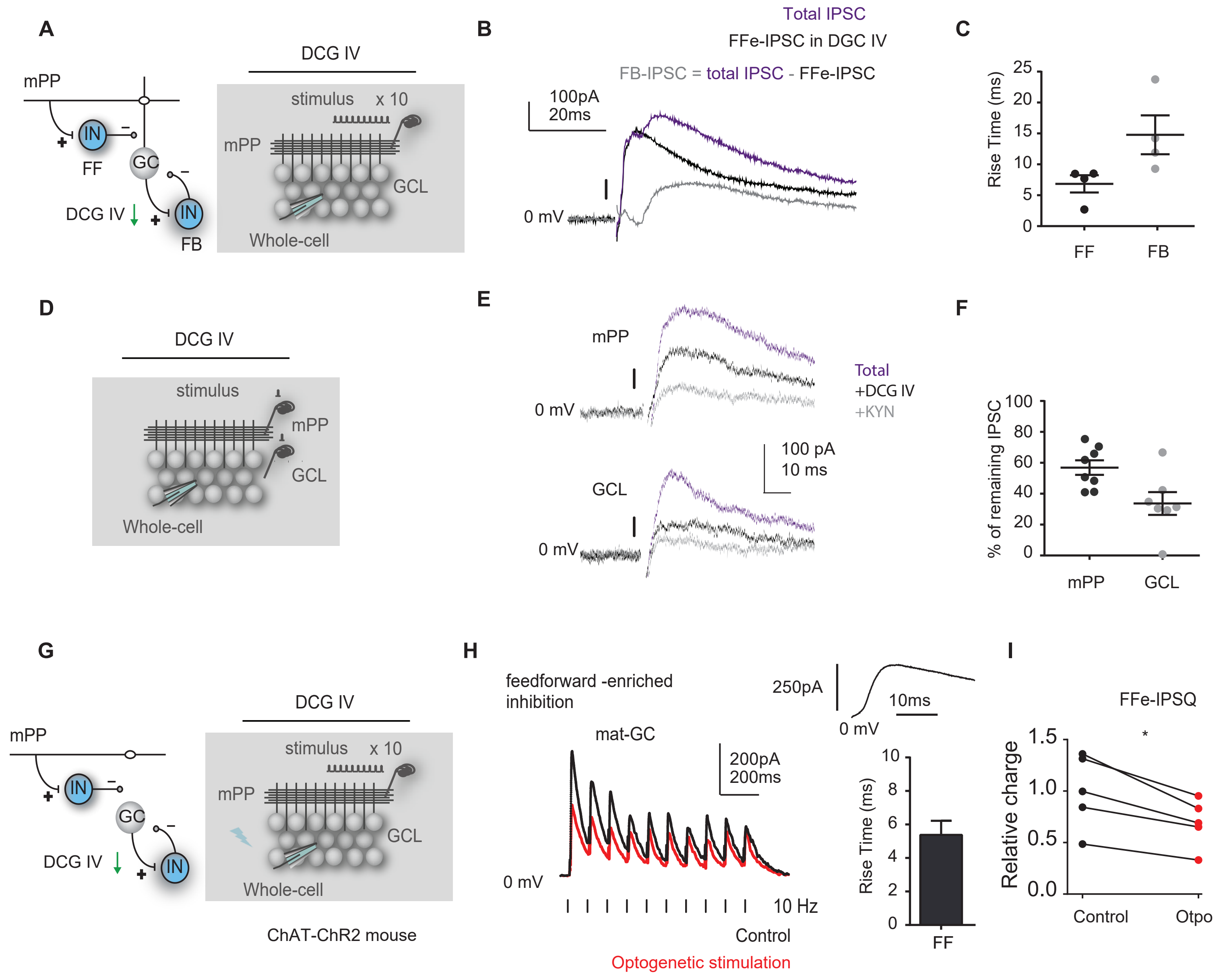
A
mPP
IN
FF
GC
IN
FB
DCG IV
DCG IV
stimulus x 10
mPP
GCL
Whole-cell
B
Total IPSC
FFe-IPSC in DGC IV
FB-IPSC = total IPSC - FFe-IPSC
100pA
20ms
0 mV
C
Rise Time (ms)
25
20
15
10
5
0
FF
FB
D
DCG IV
stimulus
mPP
GCL
Whole-cell
E
mPP
0 mV
GCL
0 mV
Total
+DCG IV
+KYN
100 pA
10 ms
F
% of remaining IPSC
100
80
60
40
20
0
mPP
GCL
G
mPP
IN
GC
IN
DCG IV
DCG IV
stimulus x 10
mPP
GCL
Whole-cell
ChAT-ChR2 mouse
H
feedforward -enriched
inhibition
mat-GC
200pA
200ms
0 mV
10 Hz
Control
Optogenetic stimulation
250pA
10ms
0 mV
Rise Time (ms)
10
8
6
4
2
0
FF
I
FFe-IPSQ
*
Relative charge
1.5
1.0
0.5
0.0
Control
Otpo

Figure S5

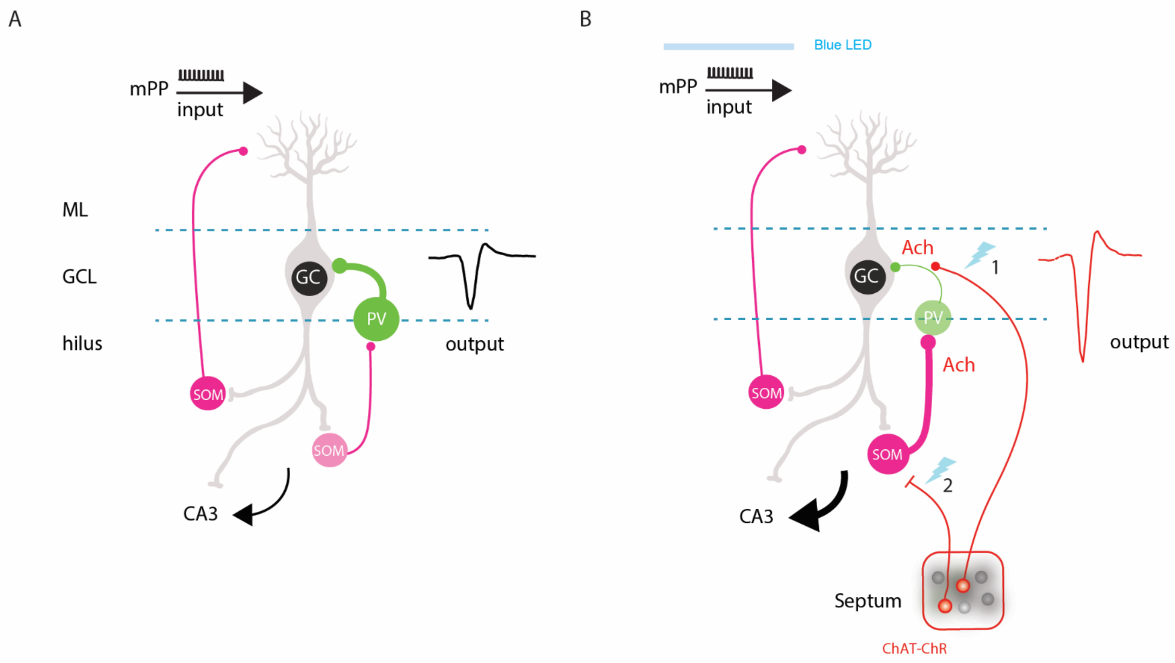
A
mPP
input
ML
GCL
hilus
GC
PV
SOM
SOM
output
CA3
B
Blue LED
mPP
input
Ach
1
GC
PV
Ach
SOM
SOM
2
output
CA3
Septum
ChAT-ChR
